# Long-term warming weakens stabilizing effects of biodiversity in aquatic ecosystems

**DOI:** 10.1101/2020.01.06.896746

**Authors:** Chun-Wei Chang, Hao Ye, Takeshi Miki, Ethan R. Deyle, Sami Souissi, Orlane Anneville, Rita Adrian, Yin-Ru Chiang, Satoshi Ichise, Michio Kumagai, Shin-ichiro S. Matsuzaki, Fuh-Kwo Shiah, Jiunn-Tzong Wu, Chih-hao Hsieh, George Sugihara

## Abstract

Despite the consensus that warming will affect biodiversity, alter physicochemical environments, and disrupt biological interactions, the relative importance of these key processes and how they interact to determine overall ecosystem function is poorly understood. Here, we analyze long-term (16∼39 years) time series data from ten aquatic ecosystems and use convergent cross mapping (CCM) to quantify the hidden causal network linking species diversity, ecosystem function, and physicochemical factors. We observe that aquatic ecosystems subject to stronger warming exhibit decreased stability (larger fluctuations in phytoplankton biomass). We further show that this effect can be attributed to a weakening of stabilizing causal pathways between biodiversity, nutrient cycling, and phytoplankton biomass. Thus, rather than thinking in terms of separate factors, a more holistic view, that causally links biodiversity and the other ecosystem components, is required to understand and predict climate impacts on the temporal stability of aquatic ecosystems.

## Introduction

Climate change has already begun to dramatically alter ecosystems (Walther *et al.* 2002), and there is growing concern about its ultimate effects on ecosystem health and resilience (Tilman *et al.* 2006; Cardinale *et al.* 2012; Fussmann *et al.* 2014). However, the mechanisms underlying ecosystem impacts are unclear, with long-term predictions that can conflict with each other. For example, some studies suggest that increased temperatures will cause decreases in biodiversity (Urrutia-Cordero *et al.* 2017; Verbeek *et al.* 2018) that will destabilize ecosystems (Hooper *et al.* 2005) and increase volatility in community biomass (Benincà *et al.* 2011); other evidence suggests that warming may actually stabilize ecosystems by altering species metabolism in ways that reduce the strength of interspecific interactions (Fussmann *et al.* 2014). The lack of consensus arises in part because of how the studies are conducted. On the one hand, experimental studies investigate the effects of climate on ecosystem properties (e.g., biodiversity and ecosystem functioning) and interactions (e.g., the effect of biodiversity on ecosystem functioning (Tilman *et al.* 2014)) in isolated single-factor style. Although tractable, this strategy is difficult to extend to large-scale manipulations of multiple interdependent processes to investigate mutual interactions and feedbacks (Hughes *et al.* 2007; Loreau 2010). On the other hand, observational studies describe the statistical relationships between ecosystem properties such as species diversity, ecosystem functioning (Grace *et al.* 2016) and stability (Ptacnik *et al.* 2008) based on linear correlative methods (Wardle 2001; Grace *et al.* 2007; Grace *et al.* 2016). These methods, however, were not designed for investigating complex interactions and feedbacks in nonlinear dynamical systems (e.g., ecosystems), and thus cannot account for interactions and ecosystem properties that change with time (Sugihara *et al.* 2012; Deyle *et al.* 2016b). Therefore, an integrated, holistic, and *dynamical* perspective is required (Chapin III *et al.* 2000) to disentangle the complex impacts of climate warming on dynamical ecosystems (Snelgrove *et al.* 2014; Dee *et al.* 2017).

We address this problem with a method specifically designed for detecting causality in nonlinear dynamical ecosystems, convergent cross mapping (CCM) (Sugihara *et al.* 2012). CCM is a causality analysis based on Takens’ theorem for dynamical systems (Takens 1981; Sauer *et al.* 1991), which infers the causal relationship among variables from their empirical time series (see Methods). With CCM, we reconstruct the causal network among diversity, ecosystem functioning, and environmental variables using long-term (16-39 years) monthly time series of phytoplankton and environmental variables from ten aquatic ecosystems spanning a wide range of geography and habitats (Fig. S1). The environmental variables consist of nutrients and water temperature (Tables S1-S3). Following previous studies, we adopt Chlorophyll-*a* concentration, a proxy for phytoplankton community biomass, as the measure for ecosystem functioning (Cardinale 2011; Lewandowska *et al.* 2016), and define ecosystem stability as the temporal stability (1 / coefficient of variation) of phytoplankton biomass (Tilman *et al.* 2006; Loreau & de Mazancourt 2013; Hautier *et al.* 2014). Here, we focus stability measure on phytoplankton biomass, as phytoplankton represent the basis of aquatic food web. We found that the systems experiencing the strongest warming exhibit the lowest ecosystem stability (Fig. 1A), which could also be observed in a global ocean dataset (Fig. 1B). We noticed that the systems Lake Mendota and Monona (Me and Mo) are leveraged, influential observations for the analysis in Fig. 1A; nevertheless, when global ocean data are included, these two systems fit reasonably well in the global pattern (Fig. 1B). That is, the general pattern that warming undermines ecosystem stability is robust in global scale. To better understand the mechanisms behind this pattern, we investigate which interaction links are associated with stability.

**Figure 1.**
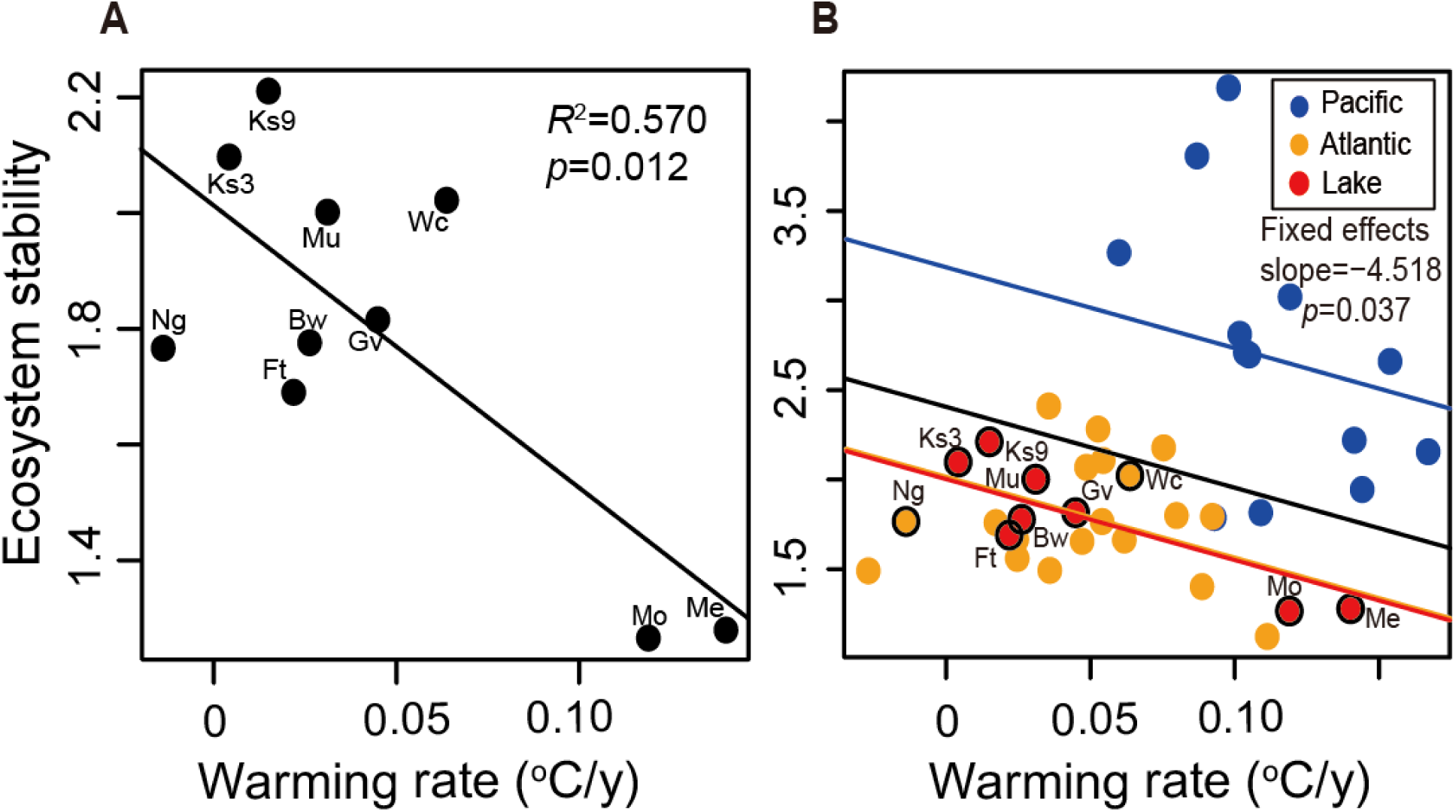
Ecosystems experiencing stronger warming are less stable (larger fluctuations in phytoplankton biomass). Ecosystem stability is measured as 1/CV of phytoplankton biomass. Here, we analyzed (A) not only the 10 monitoring sites (black circles; 24∘N∼52∘N) but also (B) globally compiled oceanic data. Ecosystem stability significantly decreased with warming rate (*p*= 0.012 and 0.037 for panel a and b, respectively). In panel (B), the black line represented the fixed effect estimated from the Generalized Mixed-effect Model (GLMM), whereas blue, orange, and red line represented the GLMM best-fit line for Pacific, Atlantic and lakes, respectively. See Table S1 for the full name of each monitoring system.

Causal interaction links inferred from CCM are denoted as X → Y for a cause X and an effect Y. A chain of connected links is a *causal pathway*, and its *linkage strength* is computed as the geometric mean of the strength associated with each of the directed links (Methods). We use the resulting networks (Supplementary Fig. 2) of links between phytoplankton species diversity, ecosystem functioning (phytoplankton biomass), and the physiochemical environment to test whether specific causal pathways are associated with ecosystem stability and to investigate how these pathways differ across the gradient of warming experienced by different systems.

**Figure 2.**
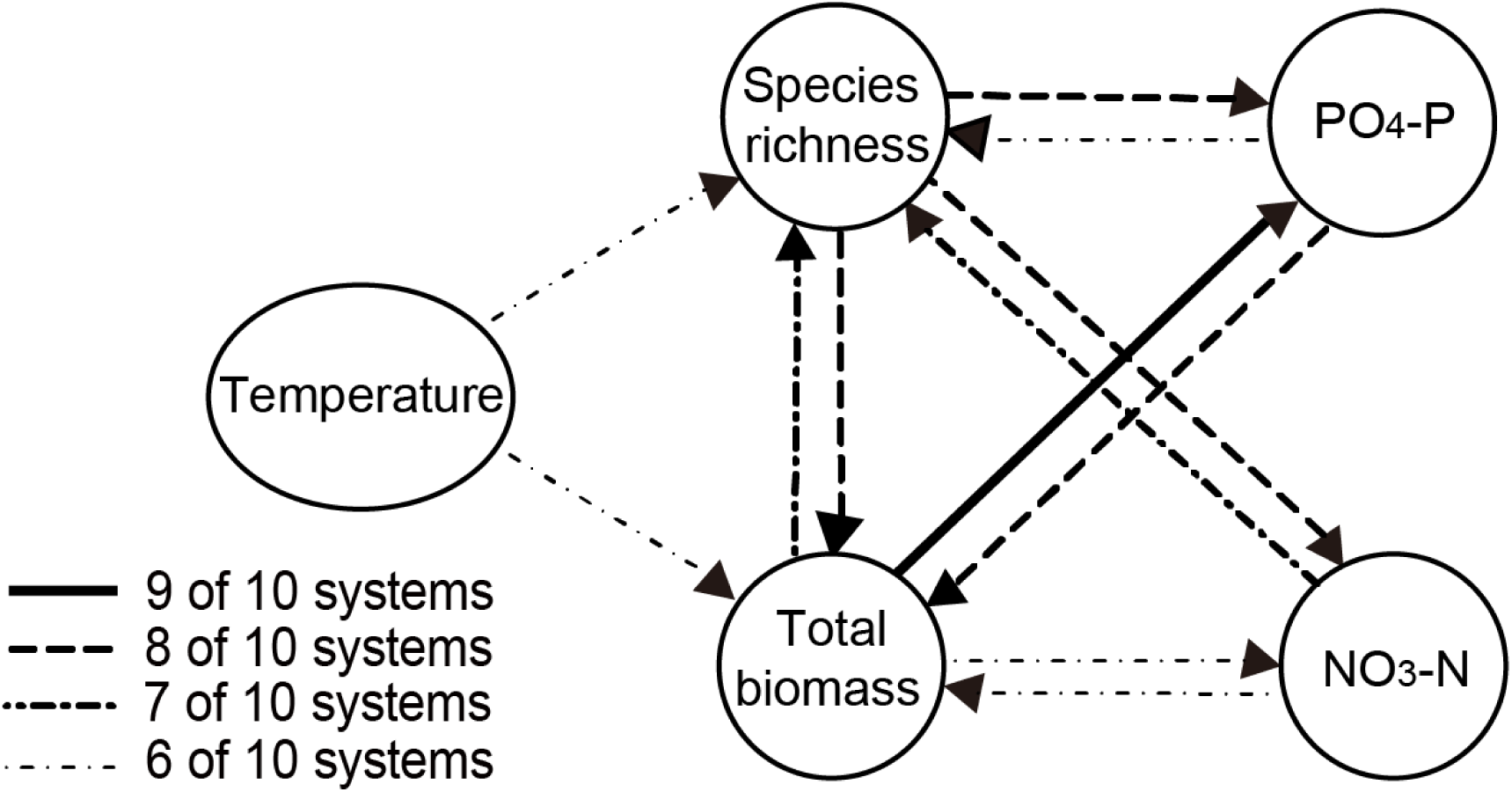
The prevalence of causal links across the ten ecosystems. Line styles denote the number of systems in which a specific CCM link is significant. Arrows indicate the direction of causality.

## Results and Discussion

Figure 2 summarizes the causal networks for the ten ecosystems, providing a general roadmap of the interactions between key ecosystem properties. Based on the reconstructed networks of individual systems (Fig. S2), we examine which ecosystem properties or interactions are associated with stability. Surprisingly, we find that ecosystem stability is not predicted by individual environmental factors (considering both their mean and variability) previously hypothesized to be relevant (Table S4), such as nutrients (Ptacnik *et al.* 2008; Hautier *et al.* 2014), water temperature (Paerl & Huisman 2008), and morphometrics (depth and area) (Hughes *et al.* 2007). Moreover, diversity indices do not show a significant positive relationship with stability (Fig. 3A, Fig. S3), even though they are usually considered an important determinant of ecosystem health and resilience (Tilman *et al.* 2006; Hooper *et al.* 2012; Tilman *et al.* 2014).

**Figure 3.**
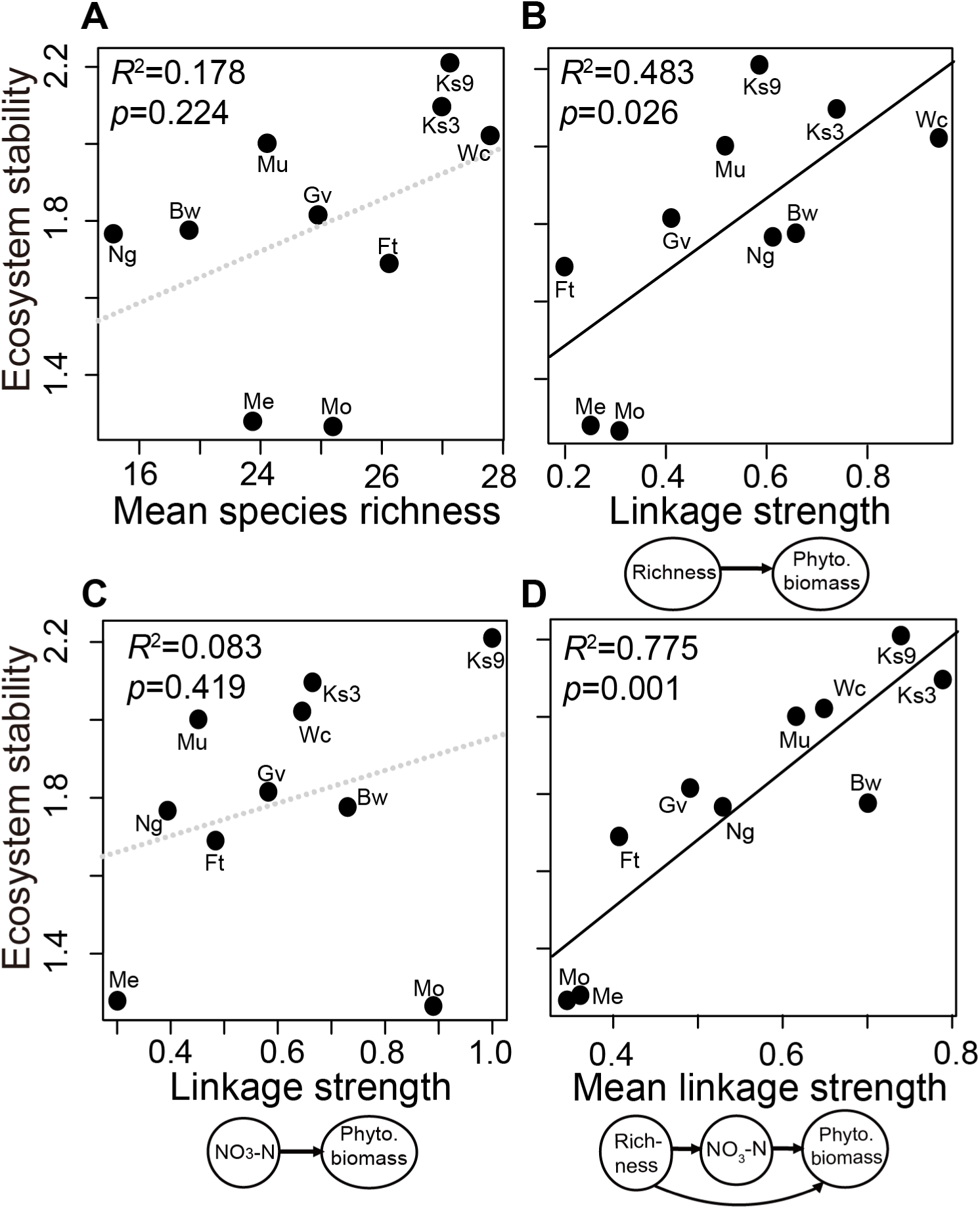
Ecosystem stability depends on BDEF and the diversity-nutrient-biomass causal pathway. (A) Mean species richness shows no significant relationship to stability. (B) BDEF strength (i.e. the effect of species richness on phytoplankton biomass) is positively associated with stability. (C) However, stability cannot be explained by the direct effect of nutrients on phytoplankton biomass. (D) Combining the nutrients and species richness into a single causal pathway further improves explanatory power (*R*^2^).

Although our analysis finds no evidence that higher biodiversity *per se* contributes to ecosystem stability, the strength of biodiversity effect on ecosystem functioning (BDEF; i.e., species richness→phytoplankton biomass), quantified in our analysis, is positively associated with stability (Fig. 3B and Table S5). While seemingly counterintuitive, there is actually a simple explanation: ecosystem properties such as species richness are not static, but are state variables of a dynamical system; stability therefore depends not only on whether the state variables have high or low values, but on the relationships between state variables. Thus, *linkage strength*, as a measure of the strength of regulatory causal pathways, is predictive of ecosystem stability, even across different systems of substantial variation in habitat type. This finding is consistent with existing hypotheses that diversity acts as a dynamical regulator of ecosystem function (Hillebrand & Matthiessen 2009). This relationship is robust to alternative measures of species diversity (Fig. S4), as well as system-specific noise (Fig. S5) and time series lengths (Fig. S6).

But what mechanisms drive this regulation effect on stability? One hypothesis is that species have different responses to environmental changes, which could help maintain community biomass at stable levels (i.e. a portfolio effect). If this is indeed the case, then species abundances should fluctuate asynchronously with high linkage strength of diversity and thus contribute to the stability of systems – which is what we observe: temporal asynchrony in species abundance driven by stronger diversity-mediated regulation is positively associated with stability (Fig. S7). Here, larger linkage strength of BDEF indicates that phytoplankton biomass responds more strongly to changing species richness, which is likely caused by low functional redundancy or high functional uniqueness in the communities (Reich *et al.* 2012). Thus, our findings of strong associations between linkage strength and stability suggest that not only diversity but functional redundancy might affect ecosystem stability. Nevertheless, we advocate advanced theoretical analysis to clarify the detailed mechanisms as well as sophisticatedly designed experiments to empirically verify the association between strengths of diversity effects on ecosystem functioning and some properties relevant to temporal stability (e.g., compensatory dynamics or response diversity).

In aquatic planktonic systems, nutrient inputs are an important driver of species turnover (Jochimsen *et al.* 2013). Indeed, the linkage strength between nutrients and species richness is a marginally significant predictor of stability (Table S5), suggesting that nutrients may influence stability. However, there is no direct correlation between nutrients and stability (Table S4), and the direct causal effect of nutrients on phytoplankton biomass is not significantly associated with stability (Fig. 3C). This indicates that species richness is a necessary intermediary for the influence of nutrients on stability.

Indeed, among all the causal pathways, the linkage strength for the pathway that includes nitrate, diversity, and biomass (more specifically, species richness→nitrate→phytoplankton biomass + species richness→phytoplankton biomass) is the best predictor of ecosystem stability (Fig. 3D). The predictive power of this pathway still significantly explained ecosystem stability, even when the influential observations (i.e., Me and Mo in Fig. 1A) are excluded from the analysis (*p*-value became 0.024). The same analysis, using phosphate instead of nitrate, produces similar results (Fig. S8). In other words, species diversity stabilizes phytoplankton biomass through regulating nutrient cycling, as suggested previously (Cardinale 2011). Typically, the ability of species diversity to stabilize phytoplankton biomass in response to nutrient fluctuations manifests as asynchronous fluctuations among different species. For example in Lake Geneva, a return to a mesotrophic state resulted in the extirpation of some phytoplankton species and the flourishing of others, with only minor subsequent changes to the net phytoplankton biomass (Anneville *et al.* 2002). This regulatory role of species diversity on nutrient fluctuations is what allows ecosystems to be less volatile. Recent studies on phytoplankton have emphasized the need to examine resource use efficiency (RUE) as an important ecosystem functioning (Ptacnik *et al.* 2008). Thus, we repeat the analyses based on variability of RUE, using a subset of our ecosystems (considering only the systems with sufficient data). We reach qualitatively similar conclusions (Fig. S9), albeit that the results are less significant due to smaller sample size. Our findings suggest that explicitly resolving the causal pathways connecting biodiversity and the other key ecosystem components offers a better understanding on the temporal stability of ecosystem functioning and can be extended to other types of ecosystems (e.g., grassland or microbial ecosystems) with better long-term monitoring.

In comparison to linear analyses (e.g. cross-correlation and structural equation modelling (SEM); Fig. S10), CCM has higher statistical power of explanation. Although both SEM and CCM considers the influences of confounding environmental variables, they include the influences of covariates in different ways. SEM has the advantage that explicitly excludes the additive confounding effects in linear systems, but it also requires correct identification of confounding variables. In contrast, CCM implicitly considers the confounding variables using lagged embeddings which does not require identifications of confounding variables, and thus can be applied in more general nonlinear dynamical systems (Ye *et al.* 2015). Moreover, in nonlinear dynamical systems where relationships between any two variables depends on other state variables (Clark & Luis 2020), linear associations will appear then disappear or change sign – so-called *mirage correlations* (Sugihara *et al.* 2012). Whereas linear correlations are ephemeral (e.g., the correlation between diversity and phytoplankton biomass changes with the time period analyzed -- see Supplementary Text and Fig. S11), CCM accounts for dynamic interactions and context-dependency and is thus able to better detect and quantify these causal links in nonlinear dynamical ecosystems.

Our analysis also gives insight into how a warming climate affects and will continue to affect ecosystem processes. Ecosystems undergoing stronger warming tend to have weaker BDEF (Fig. 4A), echoing previous experimental results in grasslands (De Boeck *et al.* 2008). Even so, just as the relationship between BDEF and stability is strengthened by including nutrients in the causal pathway (recall Fig. 3D), there is a stronger response to warming for the causal pathway that includes nutrients (Fig. 4B; *R*^2^=0.417, compared to 0.187 when only considering BDEF). This reinforces the view that long-term warming weakens the ability of the community to buffer against nutrient fluctuations, resulting in decreased stability. Moreover, the responses of these processes to warming is stronger when restricting the data to freshwater systems (*R*^2^=0.408 and *R*^2^=0.610, respectively; Fig. S12), implying different responses among biomes. More data from marine systems are needed to confirm the significance of this difference. Certainly, more datasets from other types of aquatic systems are needed to confirm our mechanistic explanation, since our key results are based on only 10 datasets.

**Figure 4.**
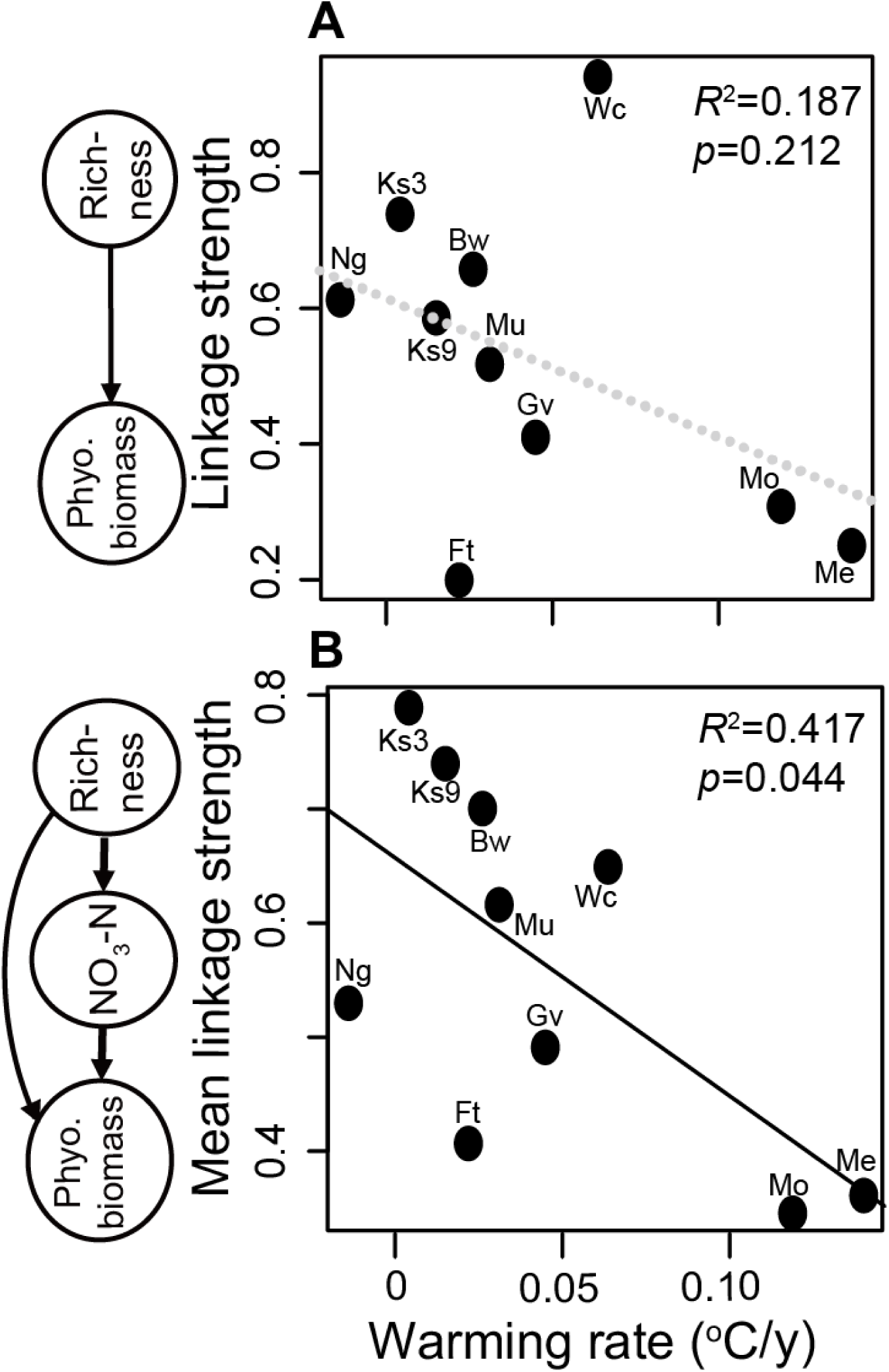
The effect of warming rate on linkage strength and ecosystem stability. The linkage strengths of BDEF (A) and the nitrate-diversity-biomass pathway (B) are weaker under stronger warming. Exclusion of the two marine systems (Wc and Ng) further improves the *R*^2^ of the relationship (Fig. S15).

Although it is recognized that warming has direct effects on individual species and their environments, its impacts on the resiliency and regulatory mechanisms of whole communities have been difficult to observe. Our results confirm the prevailing wisdom that warming makes ecosystems less resilient (Anthony *et al.* 2011; Hughes *et al.* 2019), and is therefore a key risk factor for critical shifts into undesirable states as a consequence of weakening stabilizing mechanisms. The weakening of diversity mediating causal pathways might be a consequence of preferential removal of some species with certain traits under warming (De Boeck *et al.* 2008), e.g., large body size in aquatic ecosystems (Daufresne *et al.* 2009). However, the detail mechanisms driving the preferential removal of species might be very different in different biomes. Nevertheless, the approach taken here allows us to empirically map out and quantify the ecosystem responses and causal pathways that have long been hypothesized (Anneville *et al.* 2002). This is a necessary step toward constructing quantitative indicators for ecosystem risk to enable accurate predictions and management of the future effects of climate change.

## Materials and Methods

### Data

The sampled pelagic habitats include: Lake Geneva (1974-2012), Lake Biwa (1978-2009), Lake Kasumigaura station 3 and station 9 (1977-2009), Lake Mendota (1995-2012), Lake Monona (1995-2011), Lake Müggelsee (1994-2013), Narragansett Bay (1999-2014), Western Channel (1992-2009), and Feitsui Reservoir (1986-2017) (Fig. S1). The stations 3 and 9 in Lake Kasumigaura are independent monitoring stations with distinct bottom depths (3 m and 6 m, respectively) and limnological characteristics (Table S1). Feitsui Reservoir situates at a low latitude of ∼25°N, whereas the other ecosystems are at latitudes ranging from 35° to 52° N. These ecosystems encompass a wide range of geographic regions and habitats, including both marine and freshwater systems. Detailed characteristics of these systems are listed in Tables S1-S3. For each system, data include: (i) species diversity (phytoplankton species richness); (ii) ecosystem functioning (chlorophyll-*a* concentration as a proxy for total phytoplankton biomass (Cardinale 2011)); (iii) phosphorus stock (phosphate concentration); (iv) nitrogen stock (nitrate concentration); (v) physical factors (e.g. water temperature, salinity, and irradiance). In systems with depth-resolved measurements, data are depth-integrated averages in the euphotic zone; otherwise, only surface layer measurements are used. This yielded a total of 2854 and 2790 data points for species diversity and ecosystem functioning, respectively, across the 10 sites. The available variables differ among systems (Tables S3); thus, only the variables and interactions consistently measured across ecosystems are considered in the cross-system comparisons (Figs. 3 and 4, Table S3). For example, our analysis on the effects of nutrients on ecosystem stability uses phosphate but not total phosphorous because the latter is not measured in all systems of this study. Similarly, other ecosystem functions (e.g. primary production, decomposition, respiration, etc.) are potentially meaningful metrics, but were not consistently observed across systems. We used chlorophyll-*a* concentration determined by standard spectrophotometric approach (Strickland & Parsons 1977) as a proxy for total phytoplankton biomass instead of estimation from species composition to keep the diversity and biomass data independent, following the recommendation in the literature (Ptacnik *et al.* 2008; Cardinale 2011; Ye *et al.* 2019).

All systems provide standardized phytoplankton species density data (ind./ml). Most of individuals were identified to species level under the microscope wherever possible. The counting method throughout the period of sampling is generally consistent within system, i.e., similar effort. The details of the counting method for each system are summarized in Table S2. These methods are more or less similar among systems, usually following a classic Utermöhl approach, except for Narragansett Bay using Sedgewick-Rafter technique, Lake Biwa enumerating alive plankton cells, and Feitsui Reservoir using modified Utermöhl technique because phytoplankton samples were collected and concentrated by a plankton net. We note that counting effort may vary slightly among systems. However, we have standardized the results within system before carrying out cross-system analyses (see *Convergent cross mapping analysis* below) to cope with the potential among-system difference in phytoplankton counting and identification.

### Convergent cross mapping analysis

For consistency, time series are averaged to monthly values. To make the data stationary for further time series analysis (Box *et al.* 2015), we removed the long-term linear trend and seasonality from the time series for each system. Through removing the long-term trend and seasonality, we aim to examine the variability of phytoplankton system at monthly scale, and to exclude the apparent variability simply arising from the secular trend and seasonality that may be associated with difference in latitude or ecosystem types. Seasonality is accounted for by scaling (Ye *et al.* 2015): *D*_*-mv*_(*t*_*i*_)=(*O*(*t*_*i*_)-*μ*_month *i*_)/*σ*_month *i*_, where *μ*_month *i*_ is the mean for month *i, σ*_month *i*_ is the standard deviation for month *i, O*(*t*_*i*_) is the original time series, *D*_*-mv*_(*t*_*i*_) is the deseasoned time series, and *i* = 1, 2, …, 12 corresponds to January, February, …, December, respectively. This processing method also helps to avoid the false positive results in CCM caused by ‘dynamical synchronization’ (Deyle *et al.* 2016a; Sugihara *et al.* 2017) under strong seasonality and is justified from simulations (See Supplementary texts, “<u>Justification of using de-seasoned data for CCM analyses</u>”, and Table S6). It is worth noting that, apart from seasonality, dynamical synchronization can also occur when interactions between variables are very strong (Sugihara *et al.* 2012); nevertheless, very strong interactions are of less concern here because most of interactions in real ecosystems are weak to moderate (McCann *et al.* 1998). Modelling study also indicates that CCM is robust against moderate noise from process and observational errors (BozorgMagham *et al.* 2015).

We applied convergent cross mapping (CCM) (Sugihara *et al.* 2012) to quantify causal interactions between pairs of time series, e.g. *X*(*t*) and *Y*(*t*). This method, based on Sauer, Yorke and Casdagli’s extension (Sauer *et al.* 1991) of Takens’ theorem (Takens 1981) for dynamical systems, tests for causation by measuring the extent to which the causal variable has left an imprint in the time series of the affected variable (Sugihara *et al.* 2012). That is, CCM is based on information recover (i.e., effect variables contain encoded information on causal variables), instead of predictive ability (i.e., using causal variables to predict future values of effect variables, e.g., Granger’s causality). Thus for example, if sardines are affected by temperature, it should be possible to recover past temperatures from the sardine dynamics (Sugihara *et al.* 2012). This is because temperature left its ‘footprint’ in the past history of sardines, which is preserved in the time series of sardines (Sugihara *et al.* 2012). The essential ideas of CCM are summarized in the following brief animations: tinyurl.com/EDM-intro. In this study, the embedding dimension (*E*) for each causal link (e.g. from *Y*(*t*) to *X*(*t*)) in CCM analysis was determined by testing values of *E* from 2 to 20 dimensions that optimizes the hindcast cross-mapping skill *ρ* in which *X*(*t*) is used to predict *Y*(*t*-1) (Deyle *et al.* 2016a) to prevent the overfitting of the cross-mapping between *X*(*t*) and *Y*(*t*) (Deyle *et al.* 2016a). In short, we carried out the cross-mapping between a pair of embedded variables, from [*X*(*t*), *X*(*t*-*τ*), *X*(*t*-2*τ*),…, *X*(*t*-(*E*-1)*τ*)] to [*Y*(*t*), *Y*(*t*-*τ*), *Y*(*t*-2*τ*),…, *Y*(*t*-(*E*-1)*τ*)] where *τ* is the sampling interval (one month) of time series. Although CCM detects the causations between variables in a pairwise manner, the use of lagged coordinate embedding to form state space reconstruction implicitly incorporates the influences of the critical (or confounding) environmental contexts and variables (Takens 1981; Sauer *et al.* 1991) in the dynamical system, even though these confounding variables are not explicitly specified in the embedding model (Runge *et al.* 2019).

Sugihara *et al.* (Sugihara *et al.* 2012) indicate that the link “*Y* causes *X”*, can be confirmed if points on the reconstructed attractor for *X(t)* can cross map to points on the reconstructed attractor for *Y(t)*, and that this cross-mapping converges – meaning that the cross-mapping skill *ρ*(*L*) improves with increasing library lengths (*L*; i.e., the length of training subsets from time series *X*). Because causation can occur with a lagged response (van Nes *et al.* 2015), we use the mapping with the highest cross map skill – specifically, causation between *Y*(*t*) and *X*(*t*+*k*), where *k* is a time lag equal to 0, 1, 2, or 3 months (the relevant time scales for an ecological response in this study). Convergence is determined using different library lengths (*L*) subsampled randomly from *X*(*t*), (*L*_*i*_), where *L*_*min*_= the embedding dimension, and *L*_*max*_ = the whole time series length. To test for convergence, we applied the following two statistical criteria: (1) whether there is a significant monotonic increasing trend in *ρ*(*L*) according to Kendall’s *τ* test; (2) the significance of improvement in *ρ*(*L*) (i.e. Δ*ρ*) by Fisher’s *Z* test. The latter checks whether (*ρ*(*L*_*max*_)) is significantly higher than (*ρ*(*L*_*min*_)). Convergence requires that both Kendall’s *τ* test and Fisher’s Δ*ρ Z* test are significant (i.e., max(*p*_Kendall-test_, *p*_*Z*-test_)<*α*=0.05).

As in previous studies (Sugihara *et al.* 2012; BozorgMagham *et al.* 2015) linkage strength is based on the cross-mapping skill after convergence (*ρ*(*L*_*max*_)). For linkage strengths to be comparable between systems, it is necessary to account for differences in cross-map skill resulting from differences in environmental background noise. Thus, linkage strength is scaled by dividing the cross-map *ρ* by the maximum obtained in each system, yielding standardized linkage strength (SLS) that ranges between 0 and 1, giving the relative importance of each link with respect to the strongest causal link in each system. For example, BDEF (diversity effects on ecosystem function) was defined as the SLS of species richness on phytoplankton biomass. Note that, links in causality network are constructed separately for each system; thus, we do not assume that all systems belong to the same attractor.

As a comparison with linear approaches, we computed pairwise interactions using cross-correlation and structural equation modelling (Grace 2006) (see Fig. S10).

### Linkage strength of a pathway

We quantified network pathways using the geometric mean of the SLS for all links in the pathway (analogous to loop weight (Neutel *et al.* 2002)). For example, the strength of diversity-mediated regulation, nutrient→richness→phytoplankton biomass, is computed as the geometric mean of the SLS values for nutrient→richness and richness→phytoplankton biomass. We recognize that links of causal pathways within a network are not independent (Sugihara *et al.* 2012; Anneville *et al.* 2019); nevertheless, the average interaction strength quantified for a pathway represents a reasonable approximation of joined regulatory strength along the pathways and reflects the real-world situation where links among a network are rarely independent (Levine *et al.* 2017).

### Ecosystem stability

Following previous studies (Tilman 1999; Tilman *et al.* 2006), the long-term ecosystem stability is computed as the inverse coefficient of variation (1/CV) of phytoplankton biomass. Namely, 1/CV is computed as *μ*/*σ*, where *μ* is the long-term mean calculated from the original time series; *σ* is the temporal standard deviation calculated from the detrended and deseasoned time series. Detrending and deseasoning were performed as described above, except that no variance normalization was performed in order to maintain the original variability of time series (Tilman *et al.* 2006). In this study, we focus on phytoplankton biomass, as phytoplankton represent the basis of aquatic food web.

### Evaluation of ecosystem attributes associated with stability

We use a cross-system comparison to explore associations between ecosystem stability and the linkage strength of each pathway (either an individual link or a combination of connected links). Again, the linkage strength of a pathway is either the individual link’s SLS or the geometric mean of the individual SLS values. We quantify the stabilizing effect of any pathway by how well (AIC) its linkage strength explains stability in a linear regression model. In total, the twelve links common to all ecosystems (Table S5) and their 2^12^-1 combinations were examined. In summary, we i) selected the best combination (among all candidates) using the criterion of minimizing AIC and ii) tested the statistical hypothesis only once for the selected variable and then reported the *p*-value. Therefore, we conducted only one test as if we were doing a classic stepwise regression, but not multiple tests. However, the main difference is that we keep only one independent variable (i.e., geometric mean of the individual SLS values) throughout the selection process. This makes our procedure distinctly different from step-wise multivariate regression and avoids the pitfall of over-fitting and the associated inflated Type 1 error when adding many variables in multivariate regression (Freedman & Freedman 1983; Harrell 2015).

As a comparison, we also evaluated other factors that are hypothesized to strongly influence system stability, such as diversity index (Tilman *et al.* 2006; Hooper *et al.* 2012; Tilman *et al.* 2014) and environmental variables including nutrients (Ptacnik *et al.* 2008; Lewandowska *et al.* 2016), water temperature (Paerl & Huisman 2008), and morphometrics (e.g., depth and area) (Hughes *et al.* 2007). The relationship between system stability and these factors were quantified by a linear regression model using both temporal mean and variability (CV) as explanatory variables (Table S4).

### Warming effects on linkage strength and ecosystem stability

We measured the warming rate (intensity of long-term warming) of surface water temperature using the Theil-Sen median-based trend estimator (Mohsin & Gough 2010). Then, we examined how the warming rate affects ecosystem stability and linkage strength of network pathways. First, to examine the effects of warming on ecosystem stability, we not only used the long-term time series data from the 10 monitoring stations (Fig. 1A), but also compiled oceanic data collected from two sources (Fig. 1B). The sea surface temperature data (SST) were collected with a resolution of 1*1 degree from NOAA Sea Surface Temperature V2 (Reynolds *et al.* 2002). Second, global oceanic (32°N∼59°N) chlorophyll *a* data (Chl*a*) were collected from measured Chl*a* compiled by Boyce *et al.* (Boyce *et al.* 2012). For Chl*a* data, we first averaged data into 1*1 degree as SST data. Then, all time series were averaged into a monthly basis to compare with the dataset from the 10 monitoring sites. To obtain reliable estimates of ecosystem stability from compiled Chl*a* time series, time series spanning < 10 years or containing missing values of exceeding one fourth of total time series length were excluded. In total, thirty 1*1 degree grids were analyzed. Then, a generalized mixed-effect model (GLMM) was used to examine the relationship between stability and warming rate, in which geographic region (Pacific, Atlantic and lakes) was considered a random effect, due to differences in the relationship among geographic regions.

Second, to investigate the effects of warming rate on linkage strength of network pathways, we followed a similar procedure as described above but carried out the analysis only on the 10 monitoring stations where data are available (Fig. 4).

### Sensitivity analyses with respect to time series length and ecosystem type

As a sensitivity analysis for CCM, we additionally examined the linkage strength under the same time series length specified as the minimal length (*L*_*s*_) among all the time series used in this study (i.e., around 16-year time series in Narragansett Bay) and repeated the analyses. Specifically, for any time series with the length longer than *Ls*, we compute the average linkage strength from 100 random subsamplings (library size = *Ls*) from the original time series. This randomizing procedure is performed in the **ccm** function of the R package, rEDM (assigning the arguments, random_libs=TRUE, num_samples =100) (Chang *et al.* 2017). We also repeated the analyses using only data from freshwater systems by excluding the two marine datasets (Wc and Ng). We, however, do not have sufficient data to examine exclusively marine systems.

### Computation

All analyses were done with R (ver. 3.1.2). The CCM analyses and structural equation modeling were implemented using the rEDM (Hao Ye *et al.* 2013) and lavaan (Rosseel 2012) packages, respectively. Documentation of all the analytical procedures mentioned above is provided in the Supplementary Information R script.

## Acknowledgments

We thank the participants of each long-term monitoring site. We are grateful for comments from Michio Kondoh, Masayuki Ushio, and Arndt Telschow that improved our work. Data from Lake Geneva were from SOERE OLA-IS, INRA Thonon-les-Bains, CIPEL, developed by Eco-Informatics ORE INRA Team. This work benefited from participation in the Bio-Asia FASCICLE project, the Geisha project, and the MANTEL project.

## Funding

This study was supported by the National Center for Theoretical Sciences, Foundation for the Advancement of Outstanding Scholarship, and the Ministry of Science and Technology, Taiwan (to CHH); Marie Sklodowska-Curie-Actions, H2020-MSCA-ITN_2016 (to RA); French Foundation for Research on Biodiversity and John Wesley Powell Center for Analysis and Synthesis (OA).

## Author contributions

C.W.C., C.H.H., T.M., and G.S. conceived the research idea. C.W.C. analyzed the data with help from C.H.H., E.R.D., T.M., G.S., S.S., and H.Y. O.A., R.A., Y.R.C., S.I., M.K., S.S.M., F.K.S., and J.T.W. collected data. C.W.C., C.H.H., T.M., G.S., and H.Y. wrote the manuscript with critical comments from co-authors.;

## Competing interests

The authors declare no competing interests.

## Data and materials availability

The accessibility of the time series data are provided in Table S2. Documentation of all analytical procedures is provided in the Supplementary Information R script.

## Supplementary Materials

Supplementary material for this article is available online

**Figure S1** Geographic distribution of the 10 systems.

**Figure S2** Causality network for each of the 10 ecosystems reconstructed by CCM.

**Figure S3** Relationship between rarefacted species richness and ecosystem stability.

**Figure S4** Ecosystem stability depended on strength of Shannon diversity effects on phytoplankton biomass.

**Figure S5** Ecosystem stability did not have a significant relationship with mean linkage strength of all causal links involving phytoplankton biomass.

**Figure S6** Relationship between ecosystem stability and strength of diversity-associated causal pathway was examined using a common time series length across systems in CCM.

**Figure S7** The linkage strength of diversity-mediated effects facilitates ecosystem stability through asynchronous dynamics.

**Figure S8** Ecosystem stability in relation to phosphate effects on diversity, BDEF and their combinations.

**Figure S9** Ecosystem stability based resource use efficiency (RUE) in relation to warming rate and the strength of causal pathways.

**Figure S10** Comparing performance of CCM versus cross-correlation and structural equation modeling.

**Figure S11** Mirage correlations between species richness and phytoplankton biomass in all 10 ecosystems.

**Figure S12** Relationships among BDEF, diversity-nutrient-biomass causal pathway, ecosystem stability, and warming rate in freshwater ecosystems.

**Table S1** Basic environmental information of the 10 ecosystems.

**Table S2** Data sources and information regarding long-term phytoplankton time series.

**Table S3** Accessibility of time series data among ecosystems.

**Table S4** Regression analyses between ecosystem stability versus mean and 1/CV of each of the other factors.

**Table S5** Dependence of ecosystem stability on individual causal link.

**Table S6** Justification for using de-seasoned data for CCM analyses.

