## Supplemental materials for "Long-term warming weakens stabilizing effects of biodiversity in aquatic ecosystems"

**Supplementary Materials:**

- Supplementary Text
- Tables S1-S6
- Figures. S1-S12

**Supplementary Text**

<u>Mirage correlations between biodiversity and phytoplankton biomass</u>

Ubiquitous mirage correlations between species richness and phytoplankton biomass were present in most of our studied ecosystems (Fig. S11). To determine how their relationship changed with time, moving-window analysis was used, with a 10-year window advanced by one month. Prior to analysis, seasonality was removed by subtracting long-term monthly mean. Within each window, both linear and quadratic regression analyses were performed; *l* and *q* in Fig. S11 represented the linear coefficient (simple linear regression) and quadratic coefficient (quadratic regression), respectively. Species diversity was the dependent variable, as earlier studies suggested diversity may have a unimodal relationship with productivity (i.e. communities were most diverse under an intermediate productivity level) in empirical data (Waide *et al.* 1999; Mittelbach *et al.* 2001). The model was selected based on AIC. In total, relationships between richness and total biomass were categorized into four types: 1) linear increasing ($l>0$ & $AIC_{linear}<AIC_{quadratic}$); 2) linear decreasing ($l<0$ & $AIC_{linear} <AIC_{quadratic}$); 3) unimodal ($q<0$ & $AIC_{quadratic}< AIC_{linear}$); and 4) U-shape ($q>0$ & $AIC_{quadratic}< AIC_{linear}$). Significance and type of relationship between phytoplankton diversity and biomass changed with time (Fig. S11), indicating mirage correlations.

<u>Justification of using de-seasoned data for CCM analyses</u>

Strong seasonality in time series may mask efficacy of using CCM to detect causality. Thus, prior to CCM analysis, seasonality needs to be removed. To demonstrate effectiveness of de-seasoning approaches with CCM, hypothetical time series were generated by community models with seasonality and two de-seasoning approaches were implemented and compared to analyses without de-seasoning. For the community model, we also considered two types of seasonality: additive and inseparable seasonality (i.e. seasonal-varying interaction strength), in a three-species model (*X*, *Y*, and *Z*). In both models, *Y* and *Z* were controlled by a common determinant *X* and themselves, but there was no interaction between *Y* and *Z* and no feedback influence from *Y* and *Z* to *X*. We simulated monthly dynamics of *X*, *Y*, and *Z* using the following equations:

$$\begin{aligned}
X_{t+1} &= X_t(4-3X_t)+\varphi_{1t}+0.1\varepsilon_{t,X} \\
Y_{t+1} &= Y_t(3.1-0.31\varphi_{2t}X_t-3.1Y_t)+\varphi_{1t}+0.1\varepsilon_{t,Y} \\
Z_{t+1} &= Z_t(2.12+0.2\varphi_{2t}X_t-2.12Z_t)+\varphi_{1t}+0.1\varepsilon_{t,Z}
\end{aligned} \quad (48)$$

In the additive seasonality model, $\varphi_{1t}$= 0.5*sin($2\pi t$/12), $\varphi_{2t}$=1. In the inseparable seasonality model (seasonal-varying interaction strength), $\varphi_{1t}$=0 and $\varphi_{2t}$= 0.5*sin($2\pi t$/12)+1. In both models, a minor random perturbation $\varepsilon_{t,X}, \varepsilon_{t,Y} \varepsilon_{t,Z} \sim$ *uniform*(0,1) was included. To parameterize the model, parameters were chosen based on Sugihara et al.'s example (Sugihara *et al.* 2012) which made this system exhibit chaotic behavior. Simulations were for 200 years, although only the last 30-year time series was analyzed. Simulation started at the initial value, $X_0=Y_0=Z_0=0.5$.

We compared the following de-seasoning methods: (1) no de-seasoning; (2) deducting monthly mean, $D_{-m}(t_i)=(X(t_i)-\mu_{\text{month } i})$, where $\mu_{\text{month } i}$ was the monthly mean for each of the 12 months, $i$ =1, 2, …, 12; and (3) deducting monthly mean and scaled by monthly variance, $D_{-mv}(t_i)$ =($X(t_i)$-$\mu_{\text{month } i}$)/$\sigma_{\text{month } i}$ where $\sigma_{\text{month } i}$ was the monthly standard deviation. Method (3) was used in application of CCM on salmon to remove apparent cycles (Ye *et al.* 2015); therefore, this approach was considered effective for removing seasonality but preserving dynamics. Prior to CCM, all data were normalized to zero mean and unit variance.

The model test of CCM indicated that de-seasoning was necessary when there was strong seasonality in time series data. Without de-seasoning, time series had strong synchronization, resulting in incorrect identification of causation by CCM (Table S6). Moreover, the most effective approach to remove seasonality was "deducting monthly mean and scaled by monthly standard deviation." Based on model results, this approach always identified the correct causal relationship in both types of seasonal models. In contrast, the approach that "removing only seasonal mean" cannot correctly recover the causal relationship when seasonality was inseparable. Based on our model, the de-seasoning approach through "deducting monthly mean and scaled by monthly standard deviation" successfully removed seasonality and correctly identified causation in time series data.

**Table S1** Basic environmental information of the 10 ecosystems. Data presented here are from monthly averaged values for the epilimnion for Gv, Bw, Ks3, Ks9, Me, Mo, and Mu and values for surface water for Ft, Wc, and Ng, depending on data availability.

| | **Lake Geneva** | **Lake Biwa** | **Lake Kasumigaura Sta. 3** | **Lake Kasumigaura Sta. 9** | **Lake Mendota** | **Lake Monona** | **Lake Müggelsee** | **Narragansett Bay** | **Western Channel** | **Feitsui Reservoir** |
|---|---|---|---|---|---|---|---|---|---|---|
| **Abbreviation** | **Gv** | **Bw** | **Ks3** | **Ks9** | **Me** | **Mo** | **Mu** | **Ng** | **Wc** | **Ft** |
| **Duration** | 1974-2012 | 1978-2010 | 1978-2009 | 1978-2009 | 1995-2012 | 1995-2011 | 1994-2013 | 1999-2014 | 1992-2009 | 1986-2017 |
| **Area ($km^2$)** | 580 | 670 | 220 | 220 | 39 | 13 | 7 | 380 | -- | 10 |
| **Average depth (m)** | 153 | 41 | 4 | 6 | 13 | 8 | 5 | 9 | 54 | 90 |
| **Latitude** | 46°27'N | 35°20'N | 36°12'N | 36°03'N | 43°06'N | 43°04'N | 52°26'N | 41°36'N | 50°15'N | 24°54'N |
| **Total richness** | 379 | 192 | 170 | 155 | 178 | 395 | 196 | 350 | 162 | 157 |
| **Richness range** | 7~42 | 6~34 | 1~58 | 4~57 | 9~38 | 9~52 | 9~38 | 3~25 | 9~48 | 11~42 |
| **Average richness** | 21.84 | 17.64 | 25.91 | 26.10 | 22.12 | 20.21 | 19.41 | 15.16 | 27.75 | 24.51 |
| **Chl*a* (μg/l)** | 5.52 | 4.93 | 85.14 | 53.40 | 7.56 | 9.59 | 24.10 | 6.84 | 1.51 | 3.43 |
| **Temperature (°C)** | 10.92 | 14.48 | 16.66 | 16.24 | 11.82 | 12.33 | 10.91 | 12.11 | 12.48 | 24.03 |
| **$NO_3$ (μgN/l)** | 401.06 | 125.64 | 397.21 | 133.34 | 327.10 | 135.76 | 446.14 | 31.41 | 49.15 | 417.75 |
| **$PO_4$ (μgP/l)** | 19.05 | 1.30 | 15.62 | 8.11 | 65.71 | 46.35 | 65.91 | 20.67 | 9.39 | 7.36 |

**Table S2** Data sources and information regarding long-term phytoplankton time series.

| System | Principal investigator | Contact information | Reference | Counting technique |
|---|---|---|---|---|
| Lake Geneva | Orlane Anneville | | (Anneville *et al.* 2002) | Utermöhl (Utermöhl 1958) |
| Lake Biwa | Satoshi Ichise | | (Hsieh *et al.* 2010) | Enumerate alive plankton (Kishimoto *et al.* 2013) |
| Lake Kasumigaura | Shin-ichiro Matsuzaki | http://db.cger.nies.go.jp/gem/moni-e/inter/GEMS/database/kasumi/ | (Takamura & Nakagawa 2012) | Utermöhl (Utermöhl 1958) |
| Lake Müggelsee | Rita Adrian | | (Wagner & Adrian 2011) | Utermöhl (Utermöhl 1958) |
| Lake Mendota | Stephen Carpenter | https://lter.limnology.wisc.edu/ | (Hansen & Carey 2015) | Utermöhl (Utermöhl 1958) |
| Lake Monona | Stephen Carpenter | https://lter.limnology.wisc.edu/ | (Vanni & Temte 1990) | Utermöhl (Utermöhl 1958) |
| Narragansett Bay | Tatiana Rynearson | http://www.gso.uri.edu/phytoplankton/ | (Smayda 1998) | Sedgewick-Rafter (LeGresley & McDermott 2010) |
| Western English Channel | Tim Smyth | http://www.westernchannelobservatory.org.uk | (Widdicombe *et al.* 2010) | Utermöhl (Utermöhl 1958) |
| Feitsui Reservoir | Jiunn-Tzong Wu | | (Wu & Kow 2010) | Modified Utermöhl (Wu & Kow 2010) |

**Table S3** Accessibility of time series data among ecosystems.

| | Lake Geneva | Lake Biwa | Lake Kasumi-gaura | Lake Müggelsee | Lake Mendota | Lake Monona | Narra-gansett Bay | Western Channel | Feitsui Reservoir |
|---|---|---|---|---|---|---|---|---|---|
| **Richness** | o | o | o | o | o | o | o | o | o |
| **Chl*a*** | o | o | o | o | o | o | o | o | o |
| **Water temp.** | o | o | o | o | o | o | o | o | o |
| **Light** | o | o | o | o | - | - | - | - | - |
| **Salinity** | - | - | - | - | - | - | o | o | - |
| **$NO_3$** | o | o | o | o | o | o | o | o | o |
| **$PO_4$** | o | o | o | o | o | o | o | o | o |
| **TN** | o | o | o | o | o | o | - | - | - |
| **TP** | o | o | o | o | o | o | - | - | o |

‘o’ indicates data available for analyses; ‘-’ indicates data not available.

**Table S4** Regression analyses between ecosystem stability versus mean and 1/CV of each of the other factors. The notation cor(X, Chl*a*) indicated the correlation coefficient between variable X and Chl*a* concentration. An upper case negative sign ($^-$) on the $R^2$ indicated a destabilizing effect. None of the mean and 1/CV of environmental factors significantly explained ecosystem stability. The only significant destabilizing effect was for cor($NO_3$, Chl*a*); however, its explanation power ($R^2$ and AIC) was still much weaker than causal pathway strength (Fig. 3D).

| **Variable** | **Mean** | | | | **1/CV** | | | |
|---|---|---|---|---|---|---|---|---|
| | **AIC** | $R^2$ | **rMSE** | *p* | **AIC** | $R^2$ | **rMSE** | *p* |
| Richness | 8.341 | 0.193 | 0.272 | 0.204 | 7.213 | 0.279 | 0.257 | 0.117 |
| Shannon | 6.304 | 0.342 | 0.246 | 0.076 | 9.369 | $0.105^-$ | 0.286 | 0.36 |
| Chl*a* (μg/l) | 6.764 | 0.311 | 0.251 | 0.094 | -- | -- | 0.134 | -- |
| Temperature (°C) | 10.265 | 0.022 | 0.299 | 0.685 | 10.243 | $0.024^-$ | 0.299 | 0.671 |
| TN (μgN /l) | 10.318 | 0.108 | 0.329 | 0.472 | 9.999 | 0.148 | 0.322 | 0.395 |
| $NO_3$ (μgN /l) | 10.482 | <0.001 | 0.303 | 0.974 | 9.929 | 0.054 | 0.295 | 0.519 |
| TP (μgP /l) | 10.386 | 0.056 | 0.318 | 0.574 | 10.783 | 0.008 | 0.326 | 0.837 |
| $PO_4$ (μgP/l) | 7.610 | $0.250^-$ | 0.262 | 0.141 | 10.424 | 0.006 | 0.302 | 0.832 |
| Latitude | 10.477 | 0.001 | 0.303 | 0.944 | -- | -- | -- | -- |
| Depth | 10.483 | <0.001 | 0.303 | 0.995 | -- | -- | -- | -- |
| Area | 9.714 | $0.072^-$ | 0.297 | 0.484 | -- | -- | -- | -- |
| cor(Rich, Chl*a*) | 9.088 | 0.130 | 0.282 | 0.306 | -- | -- | -- | -- |
| cor($NO_3$, Chl*a*) | 2.069 | $0.569^-$ | 0.199 | 0.012 | -- | -- | -- | -- |
| cor($PO_4$, Chl*a*) | 9.032 | 0.135 | 0.282 | 0.296 | -- | -- | -- | -- |
| cor(Temp, Chl*a*) | 7.971 | 0.222 | 0.267 | 0.169 | -- | -- | -- | -- |

**Table S5** Dependence of ecosystem stability on individual causal link. Links are ranked according to the AIC from the regressive linear model explaining ecosystem stability by standardized linkage strength of each link. Significant links ($p$ < 0.05) are underlined. An upper case negative sign (⁻) on the $R^2$ indicated a destabilizing effect.

| **Link** | **AIC** | $R^2$ | **rMSE** | ***p*-value** |
|---|---|---|---|---|
| Richness→Biomass | 3.882 | 0.483 | 0.218 | 0.026 |
| Richness→$NO_3$ | 6.422 | 0.334 | 0.247 | 0.08 |
| Temperature→Biomass | 6.577 | $0.323^-$ | 0.249 | 0.086 |
| $PO_4$→Richness | 6.861 | 0.304 | 0.253 | 0.099 |
| Biomass→Richness | 7.150 | 0.283 | 0.256 | 0.113 |
| $NO_3$→Richness | 7.180 | 0.281 | 0.257 | 0.115 |
| Biomass→$NO_3$ | 9.099 | 0.129 | 0.283 | 0.308 |
| $NO_3$→Biomass | 9.615 | 0.083 | 0.290 | 0.419 |
| Temperature→Richness | 10.134 | $0.034^-$ | 0.298 | 0.608 |
| $PO_4$→Biomass | 10.134 | $0.034^-$ | 0.298 | 0.608 |
| Richness→$PO_4$ | 10.181 | 0.030 | 0.298 | 0.634 |
| Biomass→$PO_4$ | 10.483 | $<0.001^-$ | 0.303 | 0.999 |

**Table S6** Justification for using de-seasoned data for CCM analyses. See the section of Justification of using de-seasoned data for CCM analyses in Supplementary Texts for an explanation of de-seasoning methods.

| **Seasonality** | **De-season method** | **CCM** | **Fisher's Z test** | | **Kendall's Tau** | | **Causation** | |
|---|---|---|---|---|---|---|---|---|
| | | | **delta rho** | **p** | **Tau** | **p** | **CCM converged** | **Correct identification** |
| Additive seasonality | $D_{\text{-mv}}(t_i)$ | X xmap Y | 0.360 | 0.000 | 0.198 | 0.087 | No | Yes |
| | | Y xmap X | 0.308 | 0.000 | 0.940 | 0.000 | Yes | Yes |
| | | X xmap Z | 0.367 | 0.000 | -0.315 | 0.006 | No | Yes |
| | | Z xmap X | 0.347 | 0.000 | 0.853 | 0.000 | Yes | Yes |
| | | Y xmap Z | 0.436 | 0.000 | 0.132 | 0.255 | No | Yes |
| | | Z xmap Y | 0.319 | 0.000 | -0.438 | 0.000 | No | Yes |
| Additive seasonality | $D_{\text{-m}}(t_i)$ | X xmap Y | 0.346 | 0.000 | -0.453 | 0.000 | No | Yes |
| | | Y xmap X | 0.289 | 0.000 | 0.913 | 0.000 | Yes | Yes |
| | | X xmap Z | 0.369 | 0.000 | -0.348 | 0.003 | No | Yes |
| | | Z xmap X | 0.327 | 0.000 | 0.369 | 0.001 | Yes | Yes |
| | | Y xmap Z | 0.439 | 0.000 | 0.432 | 0.000 | Yes | No |
| | | Z xmap Y | 0.323 | 0.000 | -0.477 | 0.000 | No | Yes |
| Additive seasonality | $D(t_i)$ | X xmap Y | 0.512 | 0.000 | 0.973 | 0.000 | Yes | No |
| | | Y xmap X | 0.296 | 0.000 | 0.949 | 0.000 | Yes | Yes |
| | | X xmap Z | 0.469 | 0.000 | 0.973 | 0.000 | Yes | No |
| | | Z xmap X | 0.341 | 0.000 | 0.958 | 0.000 | Yes | Yes |
| | | Y xmap Z | 0.332 | 0.000 | 0.976 | 0.000 | Yes | No |
| | | Z xmap Y | 0.359 | 0.000 | 0.973 | 0.000 | Yes | No |
| Inseparable seasonality | $D_{\text{-mv}}(t_i)$ | X xmap Y | 0.329 | 0.000 | -0.532 | 0.000 | No | Yes |
| | | Y xmap X | 0.207 | 0.002 | 0.429 | 0.000 | Yes | Yes |
| | | X xmap Z | 0.393 | 0.000 | -0.321 | 0.005 | No | Yes |
| | | Z xmap X | 0.315 | 0.000 | 0.357 | 0.002 | Yes | Yes |
| | | Y xmap Z | 0.424 | 0.000 | 0.162 | 0.162 | No | Yes |
| | | Z xmap Y | 0.326 | 0.000 | -0.297 | 0.010 | No | Yes |
| Inseparable seasonality | $D_{\text{-m}}(t_i)$ | X xmap Y | 0.347 | 0.000 | -0.318 | 0.006 | No | Yes |
| | | Y xmap X | 0.231 | 0.001 | 0.889 | 0.000 | Yes | Yes |
| | | X xmap Z | 0.399 | 0.000 | 0.685 | 0.000 | Yes | No |
| | | Z xmap X | 0.269 | 0.000 | 0.688 | 0.000 | Yes | Yes |
| | | Y xmap Z | 0.456 | 0.000 | 0.441 | 0.000 | Yes | No |
| | | Z xmap Y | 0.348 | 0.000 | 0.610 | 0.000 | Yes | No |
| Inseparable seasonality | $D(t_i)$ | X xmap Y | 0.300 | 0.000 | -0.357 | 0.002 | No | Yes |
| | | Y xmap X | 0.395 | 0.000 | 0.937 | 0.000 | Yes | Yes |

| | | | | | | |
|---|---|---|---|---|---|---|
| | X xmap Z | 0.338 | 0.000 | -0.523 | 0.000 | No | Yes |
| | Z xmap X | 0.324 | 0.000 | 0.889 | 0.000 | Yes | Yes |
| | Y xmap Z | 0.513 | 0.000 | 0.853 | 0.000 | Yes | No |
| | Z xmap Y | 0.326 | 0.000 | 0.024 | 0.844 | No | Yes |

"X xmap Y" means "X cross-map to Y".

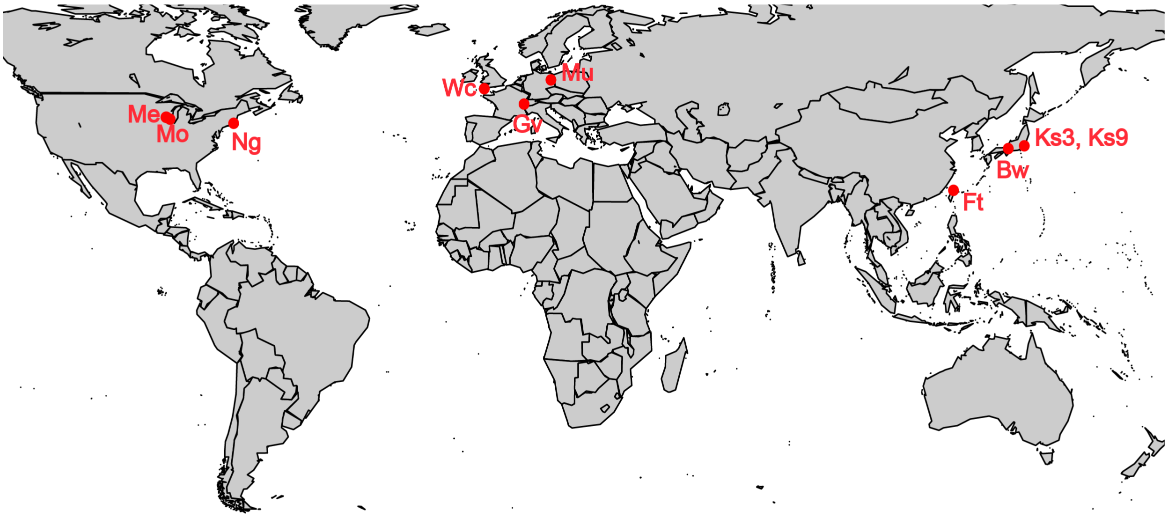

**Figure S1** Geographic distribution of the 10 systems. See Table S1 for the full name of each system.

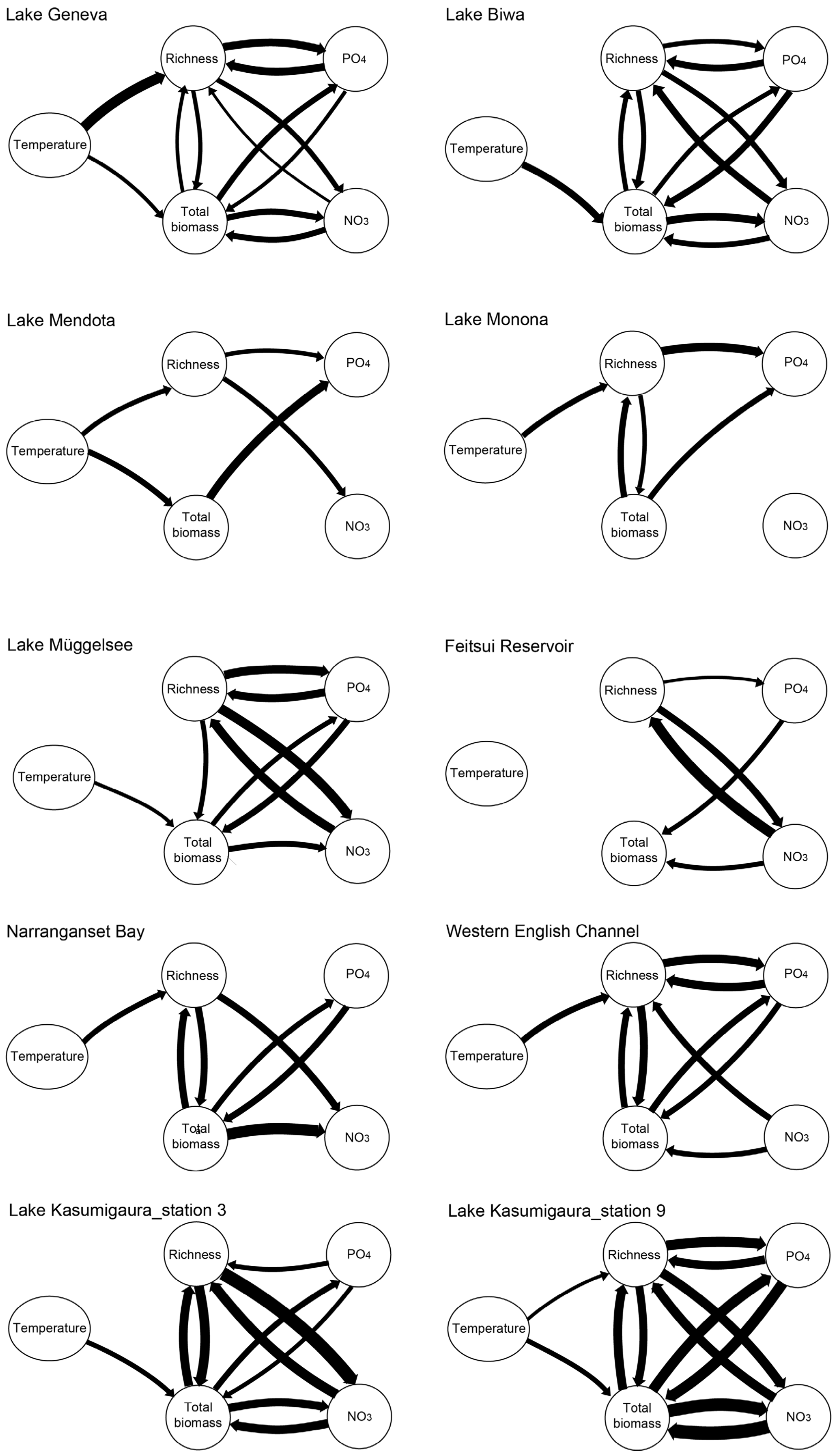


**Figure S2** Causality network for each of the 10 ecosystems reconstructed by CCM. Arrow thickness indicated linkage strength estimated by CCM.

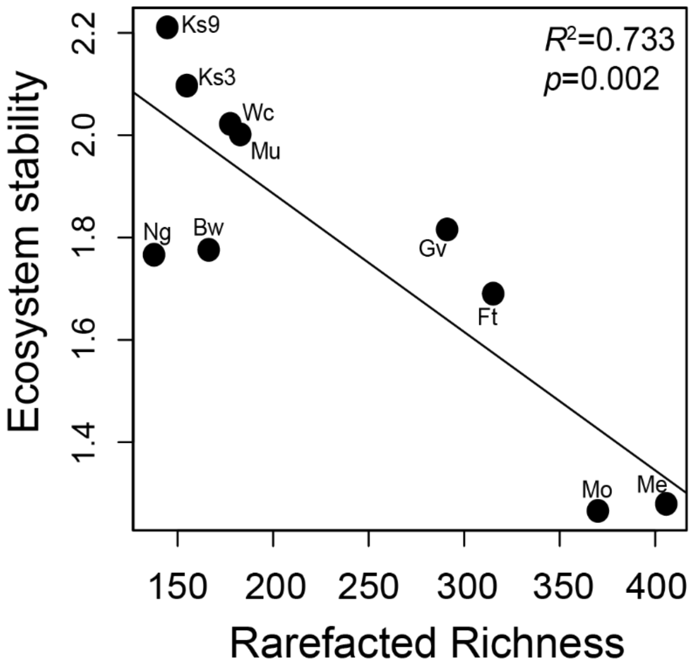


**Figure S3** Relationship between rarefacted species richness and ecosystem stability. Ecosystem stability was estimated as 1/CV of phytoplankton biomass. To ensure fair comparisons among ecosystems, species richness was standardized by Chao's rarefaction approach (Chao *et al.* 2013) which accounts for sampling effort by calculating rarefaction curves of accumulated species richness against total sampling years (approximating sampling effort). Ecosystem stability and richness had a confusing negative relationship, mainly driven by Lakes Monona (Mo) and Mendota (Me), which had the highest warming rates (Fig. 1).

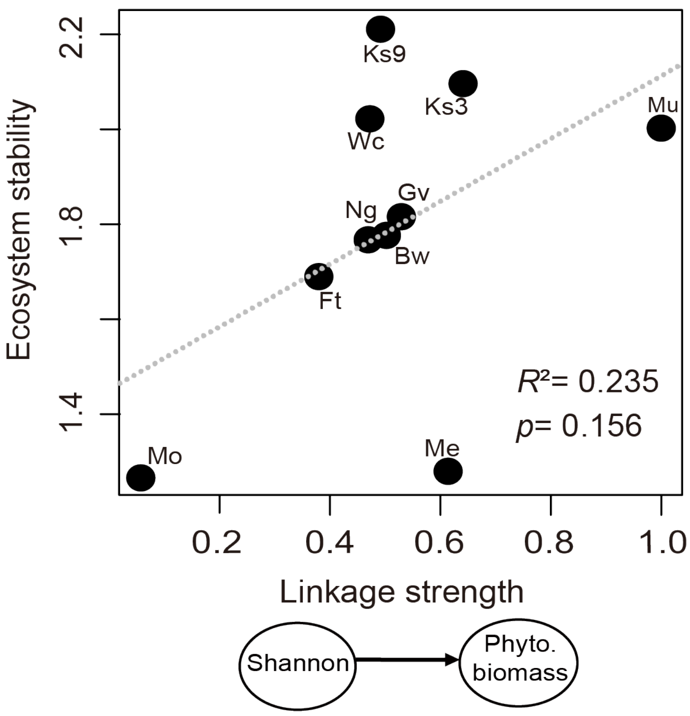


**Figure S4** Ecosystem stability depended on strength of Shannon diversity effects on phytoplankton biomass. There is a positive relationship between the linkage strength of Shannon diversity effects on phytoplankton biomass, although the relationship is not as significant as the relationship with the effect of species richness presented in Fig. 3B.

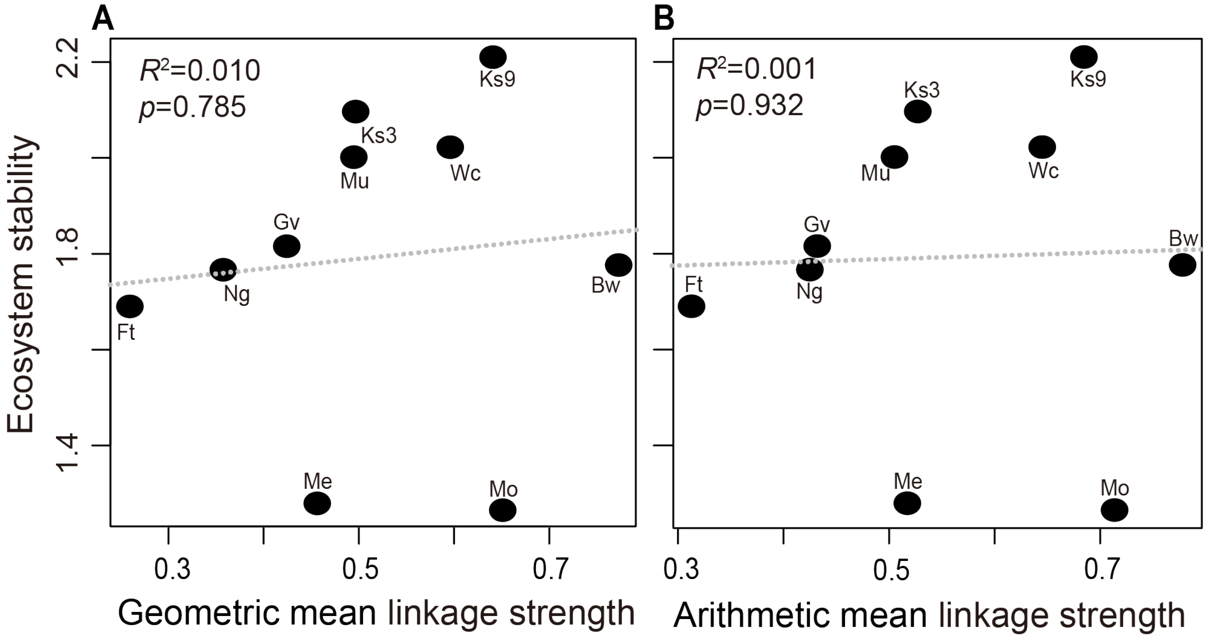


**Figure S5** Ecosystem stability did not have a significant relationship with mean linkage strength of all causal links involving phytoplankton biomass. One may suspect that our finding of positive association between stability and linkage strength (Fig. 3B, C, and D) was a statistical artifact: if the CV of phytoplankton biomass was solely driven by noise, the high predictability by CCM (strong linkage strength) simply implied low variability of biomass (high stability). To exclude this possibility, we computed (A) geometric and (B) arithmetic mean linkage strength of all causal links involving phytoplankton biomass (by nitrate, phosphate, temperature and species richness). These measurements and all other individual effects involving phytoplankton biomass (Table S5) were expected to be highly correlated with stability, provided that stability and predictability were both determined solely by noise. However, our analysis excluded this possibility, as none of the causal links significantly explained stability, except for diversity-mediated linkages. In addition, some causal links without directly involving phytoplankton biomass, e.g., species richness→nitrate, were able to explain stability (Table S5). In addition, standardized linkage strength for cross-system analyses were used to remove potential bias among systems (Methods).

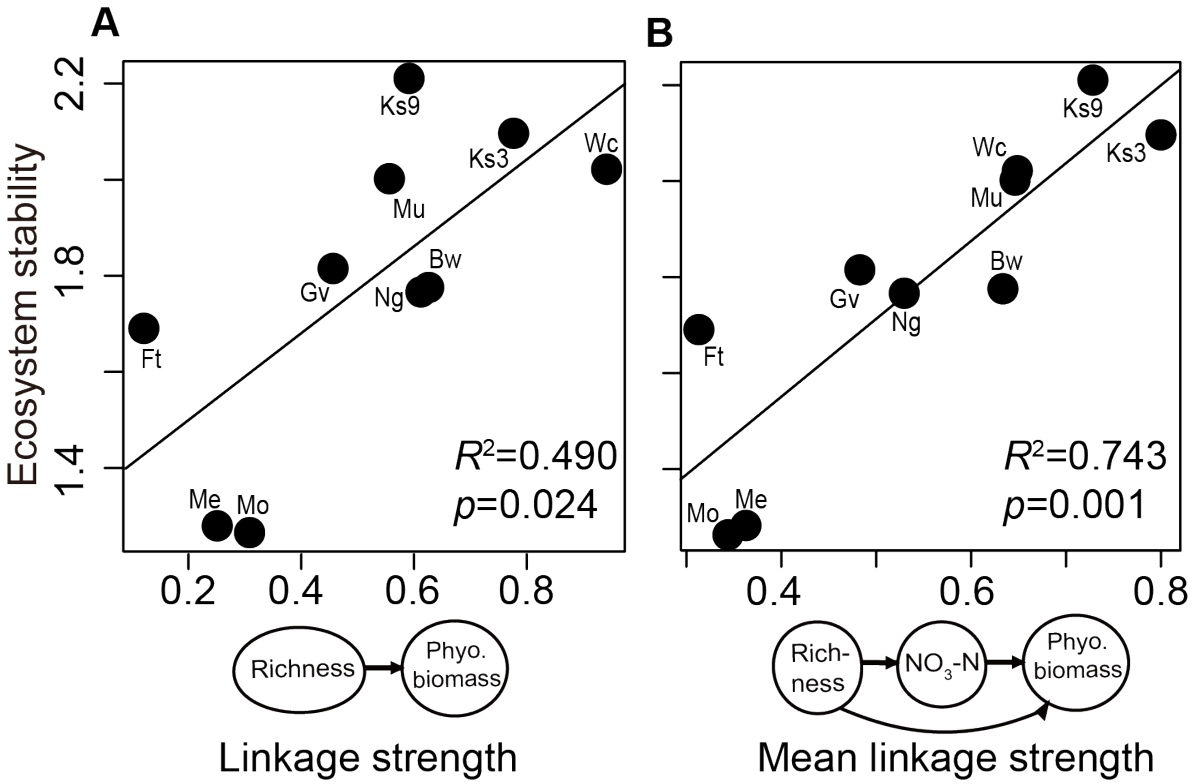


**Figure S6** Relationship between ecosystem stability and strength of diversity-associated causal pathway was examined using a common time series length across systems in CCM. Here, linkage strength of BDEF and diversity associated causal pathways were estimated under the same randomized subsampling CCM library size, as explained in Methods (Sensitivity analyses with respect to time series length and ecosystem type). Results were qualitatively consistent with those in Fig. 3. Thus, the positive relationship between the linkage strength and ecosystem stability was robust to varying length of time series observed in various systems.

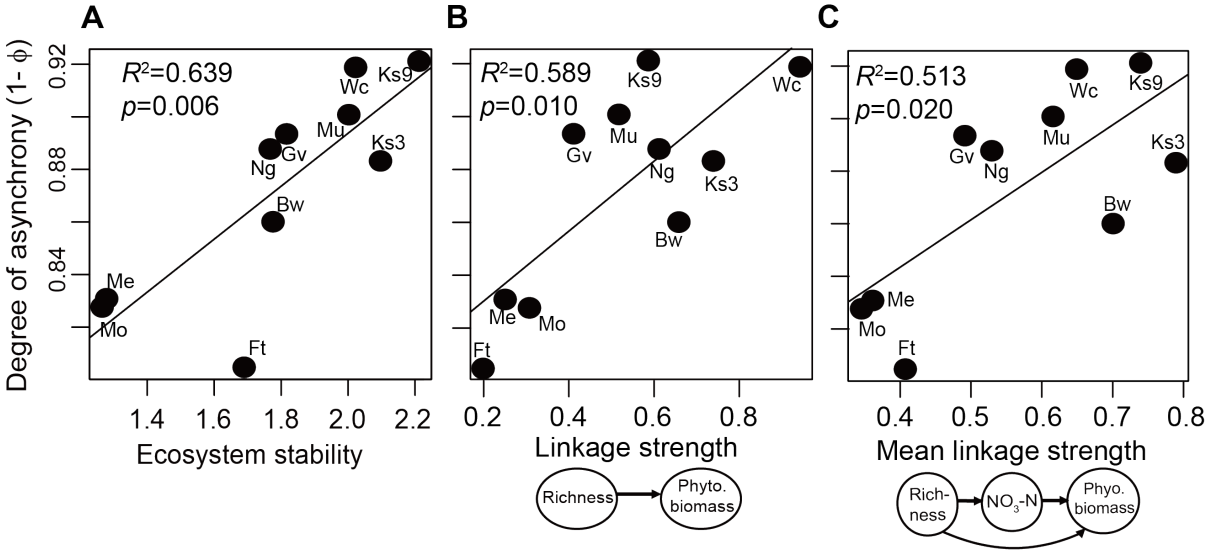


**Figure S7** The linkage strength of diversity-mediated effects facilitates ecosystem stability through asynchronous dynamics. (A) The asynchrony index calculated from dominant species had a positive relationship with ecosystem stability. The asynchrony index (Loreau & de Mazancourt 2008) (1- $\phi$) is calculated by one minus the standardized variance ratio $\phi$ and the dominant species is defined as those which attain 5% or greater biomass at any time during the study. Number of dominant species varied from 29.1 to 100% of species richness across samplings. Degree of asynchrony increased with linkage strength of species richness effects on phytoplankton biomass (B) and mean linkage strength of the following pathway: species richness $\rightarrow$ phytoplankton biomass + species richness $\rightarrow$ nitrate $\rightarrow$ phytoplankton biomass (C). We inferred that species in a community under stronger diversity-mediated regulations had stronger differential responses to environmental fluctuations (i.e. higher asynchrony); as such, the system was more stable. These results were robust to the definition of dominant species. For example, when defined as >10% total biomass, relationships were still significant, with $R^2$=0.65, 0.61, and 0.53 for Panels (A), (B), and (C), respectively.

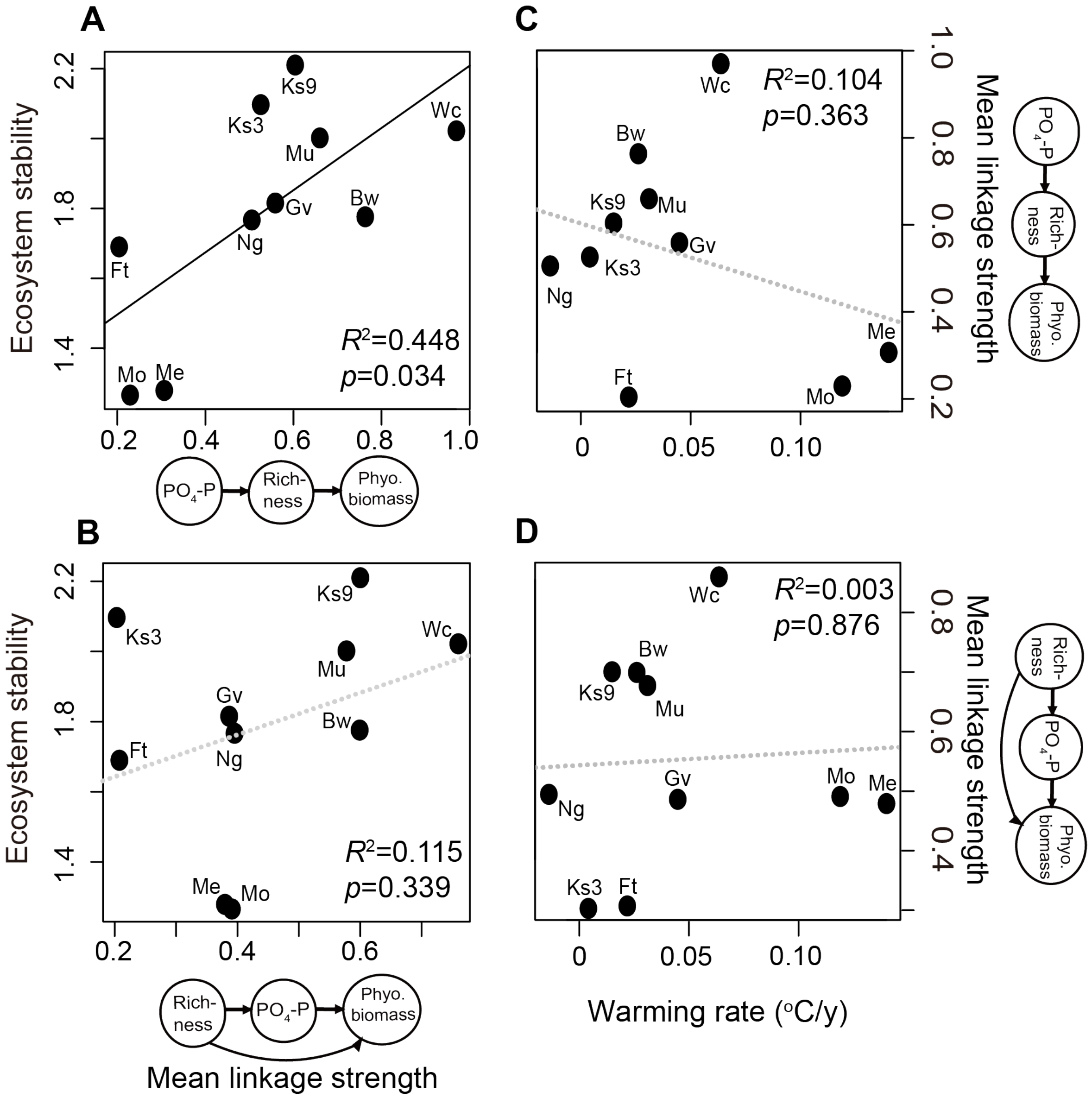


**Figure S8** Ecosystem stability in relation to phosphate effects on diversity, BDEF and their combinations. Stability (1/CV of phytoplankton biomass) significantly increased with the strength of causal pathway: phosphate effects on diversity combination with BDEF (A), similar to nitrate results (Fig. 3). However, unlike nitrate results, involving diversity effects on phosphate did not further improve explanation power (B). We inferred that both nitrate and phosphate involve stabilizing mechanisms on ecosystem functioning, although their ways of regulation may be different, driven by either diversity → nutrient ($NO_3$) or nutrient ($PO_4$) → diversity. Considering warming effects on linkage strength as presented in Fig. 4, (C) strength of the phosphate effects on species diversity combined with BDEF (D) and diversity effects on phosphate had no or negative relationship with warming condition, although results were both not significant. Perhaps the relationship between warming and its linkage strength were not well described by a linear model.

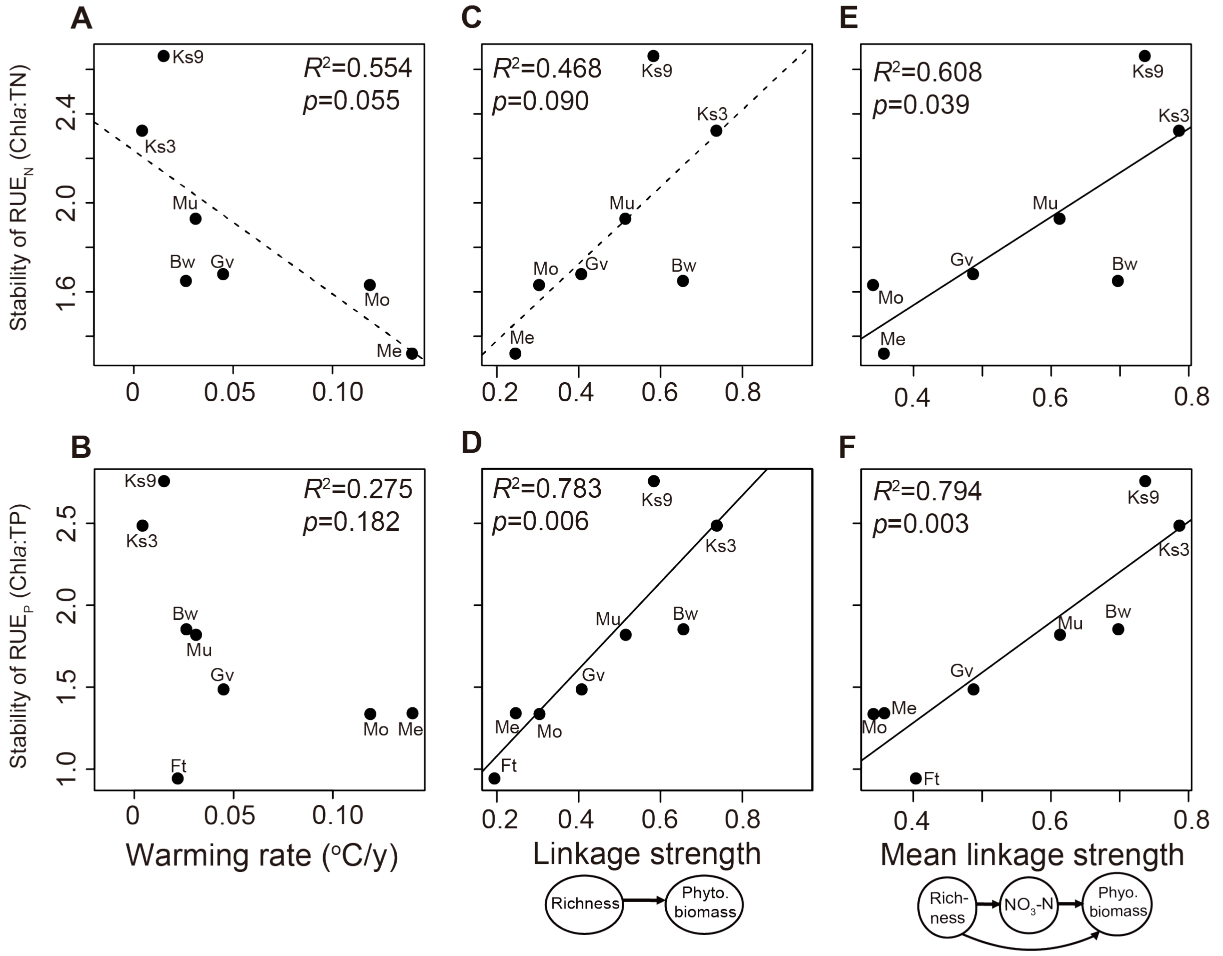


**Figure S9** Ecosystem stability based resource use efficiency (RUE) in relation to warming rate and the strength of causal pathways. RUE is defined as the ratio between Chl*a* and total nitrogen ($RUE_N$) or the ratio between Chl*a* and total phosphorous ($RUE_P$) (Ptacnik *et al.* 2008). Then, stability of RUE is calculated as 1/CV of RUE. Similar as the warming effects on Chl*a* stability as presented in Fig. 1A, the stability of (A) $RUE_N$ and (B) $RUE_P$ shows a negative association with the warming rate, albeit that the relationships are not significant due to a smaller sample size (lack of TN and TP data in some systems; Table S3). Nevertheless, both RUE stabilities increase with the strength of (C-D) BDEF and (E-F) the causal pathways (consistent with the results in Fig. 3).

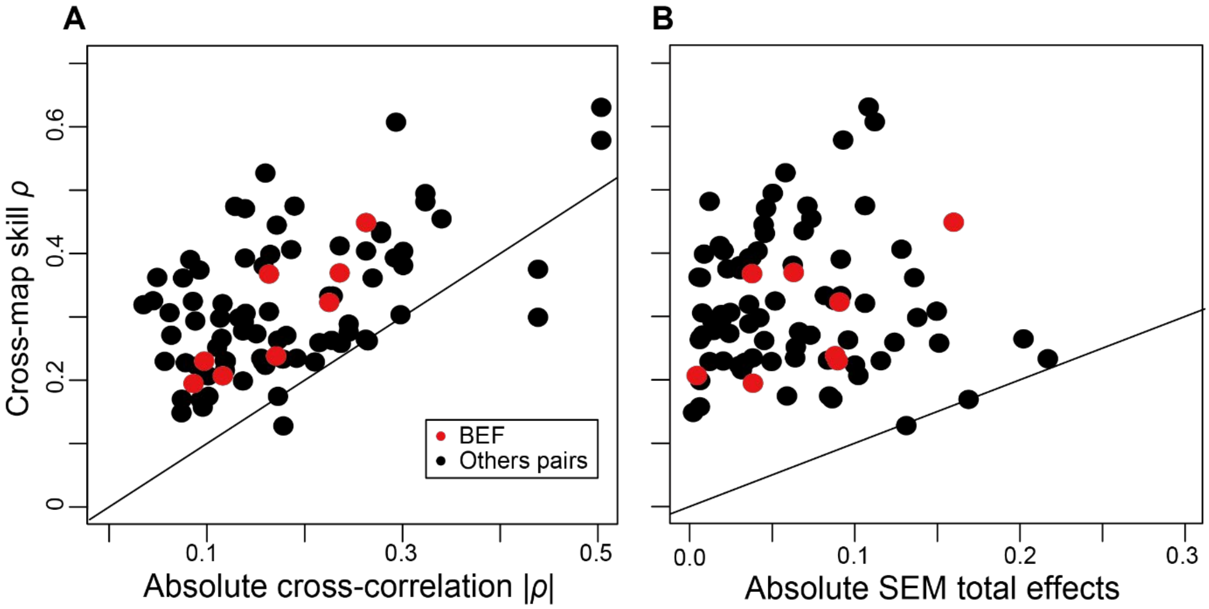


**Figure S10** Comparing performance of CCM versus cross-correlation and structural equation modeling. For each variable pair with significant CCM causation, performance of CCM (cross-map skill) was compared to: (A) best cross correlation; and (B) total effect calculated from structural equation modeling (SEM) for the variable pair. Red dots highlighted linkage strength of species richness to phytoplankton biomass (i.e., BDEF). In a majority of variable pairs and all BDEF pairs, cross-mapping skills of CCM were higher than the absolute value of cross correlation coefficients and SEM total effects. A solid line indicates the 1:1 line. To ensure valid comparisons, cross correlation also allows a three-month lag response as in CCM and the lag with the best correlation coefficient was selected. For SEM modeling, a non-recursive (bidirectional) model was analyzed to estimate feedback between variables. This analysis measured feedbacks by specifying the effect arrow pointed from a set of cause variables to another set of effect variables with a one-month time lag (i.e. multi-time step analysis) (Grace 2006). Herein, we analyzed the SEM model according to the same causality network identified in CCM.

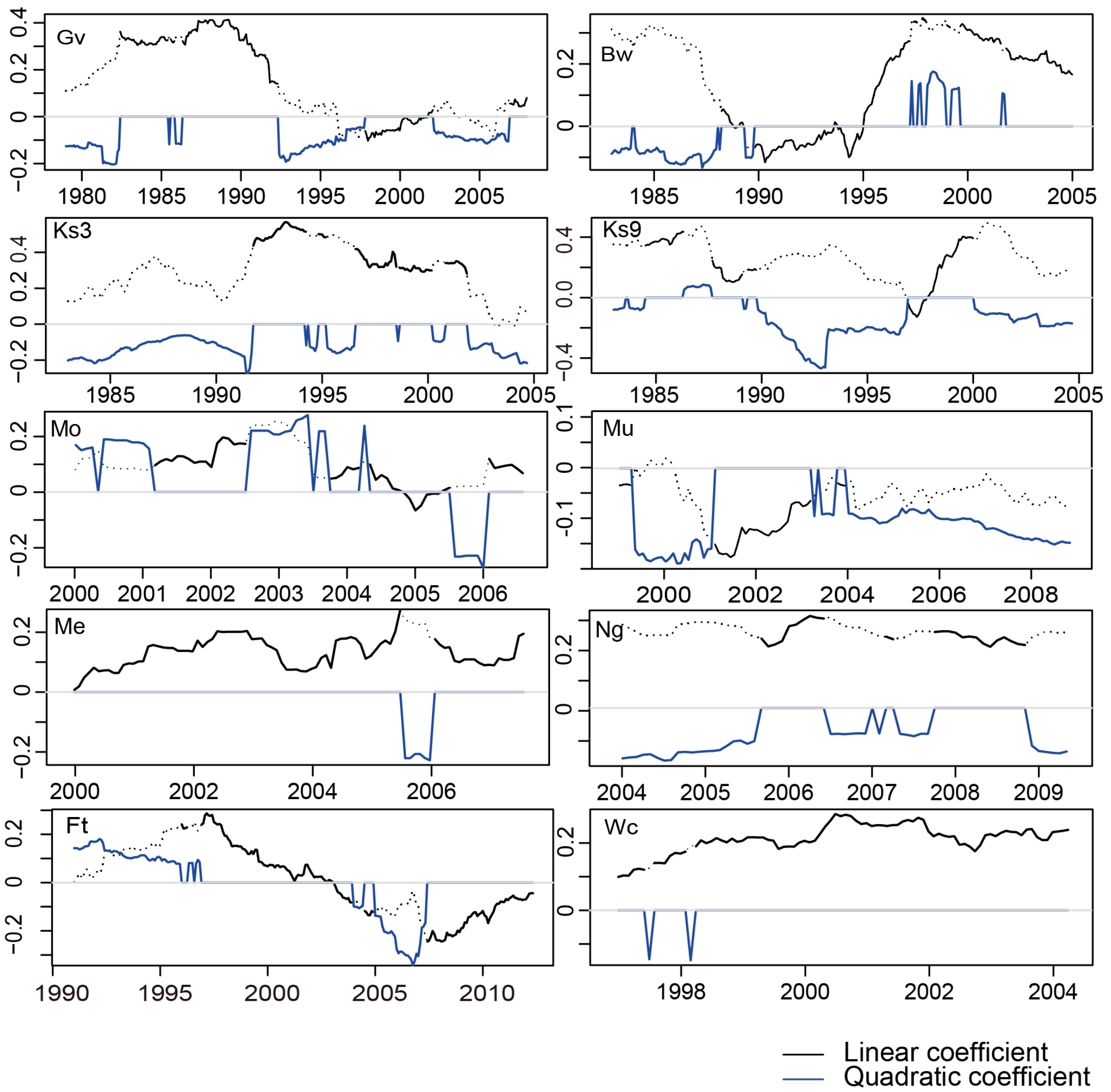


**Figure S11** Mirage correlations between species richness and phytoplankton biomass in all 10 ecosystems. The model was selected based on AIC. The black line indicated dynamics of linear regression coefficient *l*. However, if the quadratic model was selected ($AIC_{quadratic} < AIC_{linear}$), quadratic coefficient *q* was shown instead using blue line and solid black line (linear regression coefficient) was changed to dash line.

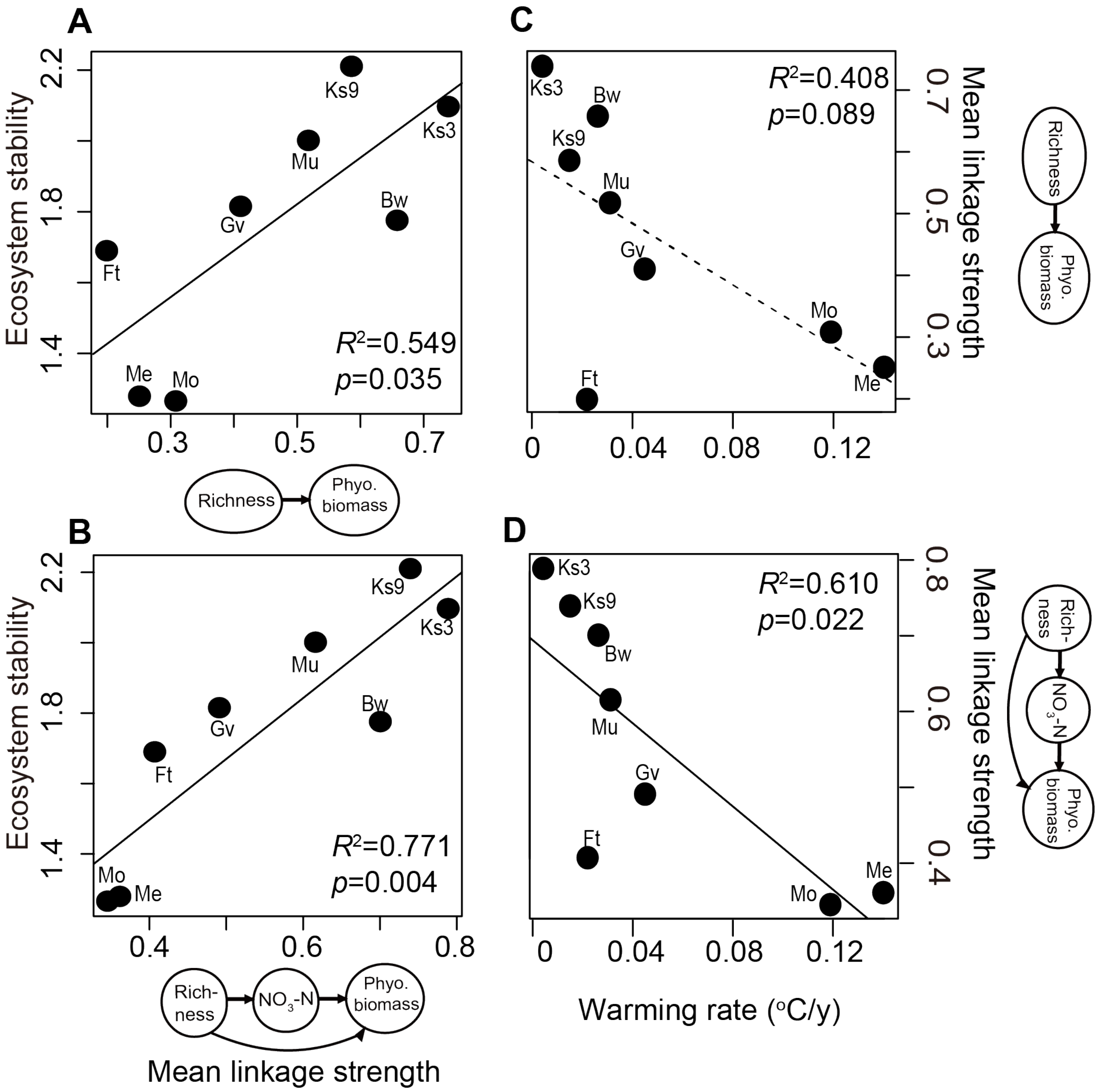


**Figure S12** Relationships among BDEF, diversity-nutrient-biomass causal pathway, ecosystem stability, and warming rate in freshwater ecosystems. In this analysis, marine datasets (Wc and Ng) were excluded. Similar to analyses presented in Fig. 3B and 3D, ecosystem stability had a significant positive relationship with the strength of (A) BDEF and (B) causal pathway including nutrients, biomass, and species richness, when considering only the freshwater systems. Moreover, strength of both (C) BDEF and (D) the causal pathway were also weaker under a rapid warming condition, similar to analyses presented in Fig. 4. The $R^2$ was higher and more significant in this analysis, excluding marine datasets, especially for the negative relationship between strength of BDEF and warming rate. An improved $R^2$ implied that causal pathways may respond to warming differently between marine and freshwater ecosystems. However, more long-term monitoring for marine systems is needed to clearly detect a differential response.
